# Time-series transcriptomics of grapevine deacclimation reveals chilling-dependent genetic responses to temperature increase during dormancy

**DOI:** 10.1101/2025.08.08.669328

**Authors:** Hongrui Wang, Jason P. Londo

**Affiliations:** Ecophysiologie et Génomique Fonctionnelle de la Vigne (EGFV), INRAE, ISVV, Villenave d’Ornon, France; School of Integrative Plant Science, Horticulture section, Cornell University-Cornell AgriTech, Geneva, NY, USA

**Keywords:** bud dormancy, deacclimation, cold hardiness, chilling accumulation, temperature response, RNA-seq, *Vitis vinifera*

## Abstract

During winter, grapevine (*Vitis vinifera*) bud dormancy and cold hardiness are regulated by complex interactions between chilling accumulation and temperature cues. However, the molecular mechanisms underlying physiological transitions during winter remain poorly understood. In this study, we performed time-series RNA-seq on ‘Cabernet Sauvignon’ dormant buds with varying chilling accumulation, followed by warm temperature exposure. Using weighted gene co-expression network analysis, empirical modeling, and a novel calculation of molecular temperature response rate, we identified gene expression patterns responsive to temperature alone, chilling alone, and their interaction. Temperature-responsive genes showed rapid, chilling-independent activation and were primarily associated with metabolism, environmental sensing, and auxin signaling. Chilling-responsive genes were enriched for functions of chromatin remodeling and heat shock protein pathways, suggesting progressive cellular reprogramming under field conditions. Interaction-responsive genes, including those involved in ABA/auxin metabolism and cell wall modification, seem to function in both dormancy progression and deacclimation. These findings provide a mechanistic framework for how chilling and temperature synergistically regulate dormancy transitions in grapevine, which enhances the understanding of temperature sensing and response and the chilling-mediate dormancy progression underlying grapevine dormant season physiology.

## Introduction

Driven largely by human activities, climate change is marked by persistent shifts in temperature and precipitation patterns, alongside a rise in the frequency and intensity of extreme weather events (Santos *et al*., 2020). These changes are disrupting global forest distributions and impacting the long-term sustainability of agricultural systems worldwide (Raza *et al*., 2019; Salama *et al*., 2021). Perennial crops are especially vulnerable to these climatic stresses due to their long life cycles and high re-establishment costs, which limits the feasibility of rapid adaptation strategies such as cultivar replacement. In the growing season, heat stress and drought stress impair physiological function and metabolite biosynthesis, which ultimately lowers the quality of fruits and derived products like wine and cider (AghaKouchak *et al*., 2020; van Leeuwen *et al*., 2024). During the dormant season, warmer average winter temperatures can increase the risk of cold damage by disturbing the process of cold acclimation, particularly when followed by extreme freezing events (Kovaleski, 2024). This is particularly true for the cultivation of domesticated grapevines (*Vitis vinifera*) in cool or cold climate regions in North America and Eurasia, where the lowest temperatures occasionally drop below their maximum cold hardiness (Zabadal *et al*., 2007). In addition, more frequent spring frost damage, a global climatic concern, can be associated with high crop loss in grapevine as the damage often occurs in primary buds, which are the most fruitful (Wang & Dami, 2020; Persico *et al*., 2021; Poni *et al*., 2022). A deeper understanding of grapevine dormant season physiology is essential for safeguarding the sustainability of current production and supporting the continued expansion of the grape and wine industries under climate change.

Grapevines and other perennial species undergo specific physiological changes to enter and maintain dormancy throughout winter, while simultaneously developing cold hardiness to withstand subfreezing temperatures. Dormancy, characterized by the cessation of visible growth, is generally classified into three categories: paradormancy, endodormancy, and ecodormancy (Lang, 1987). Paradormancy refers to the suppression of growth caused by physiological factors, primarily auxin-mediated apical dominance, during the growing season, such as the inhibition of lateral bud outgrowth by the shoot apex (Fan *et al*., 2023). At the end of the growing season, grapevine buds transition from paradormancy to endodormancy in response to shortening daylength and decreasing temperatures (Fennell, 2004; Grant *et al*., 2013). During endodormancy, growth is suppressed and buds have reduced sensitivity to environmental cues, a mechanism that protects it from responding prematurely to fluctuating conditions (Yang *et al*., 2021). Concurrent with the onset of endodormancy, grapevines transition from a cold-sensitive to a cold-hardy state through the process of cold acclimation (Gusta *et al*., 2005). To resume growth in spring, grapevines must transition from endodormancy to ecodormancy, where environmental sensitivity is restored but growth remains suppressed due to the ambient low temperature. When ecodormant buds are exposed to warm temperatures, they begin to lose cold hardiness through a temperature-dependent process known as deacclimation (Pagter & Arora, 2013; Wisniewski *et al*., 2018; Kovaleski *et al*., 2018). The transition from endodormancy to ecodormancy is thought to be triggered by the accumulation of chilling, the cumulative exposure to temperatures within a specific range during the dormant season (Dokoozlian, 1999; Campoy *et al*., 2011; Wang *et al*., 2020; Kovaleski, 2022). Various models have been developed to quantify chilling accumulation, with widely used approaches such as the North Carolina model (Shaltout & Unrath, 1983), the Utah model (Richardson *et al*., 1974), and the Dynamic Model (Fishman *et al*., 1987). These models generally consider temperatures between 0°C and 15°C as effective for chilling, though they differ in their optimal temperature ranges and in how they weight the efficiency of chilling accumulation. Recently, a new model expanded the range of chilling-effective temperatures to include freezing conditions and incorporated temperature fluctuations as an additional factor, resulting in improved chilling estimates that better correlate with physiological changes (North *et al*., 2024).

Building on this foundational understanding of dormant season physiology, studies have begun to uncover the underlying mechanisms that regulate the changes in cold hardiness and dormancy status. In grapevines, cold acclimation is typically divided into three sequential phases, initiated by shortening daylength, followed by exposure to low above-freezing temperatures (4-10°C), and finally by subfreezing temperatures (<0°C) (Fennell, 2004; Zabadal *et al*., 2007; Grant *et al*., 2013). Daily temperature fluctuations are also critical for initiating cold acclimation (Wang *et al*., 2024b; North *et al*., 2024). During cold acclimation, cold hardiness is progressively enhanced through a complex interplay of genetic regulation, metabolic adjustments, and physiological adaptations. Key genetic responses include the activation of hormone signaling pathways, such as abscisic acid (ABA), and in many species, the inducers of CBF Expression– C-Repeat Binding Factor/DRE Binding Factor–Cold Regulated Genes (*ICE-CBF/DREB1-COR*) cascade are activated, which together promote the production of functional proteins and metabolites. These molecular responses drive downstream physiological and structural changes, including cell dehydration, membrane lipid remodeling, and increased supercooling capacity, all of which contribute to enhanced cold hardiness (Noriega & Pérez, 2017; Londo *et al*., 2018; Liu *et al*., 2019; Chai *et al*., 2019; Ritonga & Chen, 2020; Ouyang *et al*., 2021; Hwarari *et al*., 2022; Noriega *et al*., 2024; Ralser *et al*., 2024).

Grapevine deacclimation is closely linked to chilling accumulation: as chilling exposure increases, grapevine deacclimation rates (*k*_*deacc*_) increase, however, substantial variation in chilling requirements and maximum deacclimation rates has been observed among *Vitis* cultivars with diverse genetic backgrounds (Londo & Johnson, 2014; Londo & Kovaleski, 2025). At the molecular level, the deacclimation of grapevine has been found to be correlated with a plant hormone metabolism and signaling network that involves ABA degradation and signaling, ethylene biosynthesis and signaling, jasmonate and gibberellin biosynthesis, auxin signaling and cytokinin degradation along with the activation of growth-related pathways that includes cell wall modification, ribosome biogenesis and photosynthesis protein biosynthesis (Kovaleski & Londo, 2019; De Rosa *et al*., 2025; Wang *et al*., 2025).

While the current understanding of grapevine dormant season physiology largely centers on chilling accumulation as the driver of dormancy transitions, the underlying mechanisms of chilling perception and response remain poorly understood. It is widely believed that the chilling-mediated transition from endormancy to ecodormancy is a consequence of plant hormone interaction and their downstream signaling (Atkinson *et al*., 2013; Brunner *et al*., 2014; Beauvieux *et al*., 2018; Yang *et al*., 2021; Penfield *et al*., 2021). The onset of endodormancy is characterized by elevated levels of ABA and reduced levels of gibberellins (GAs), whereas endodormancy release is associated with an increase in gibberellin concentrations (Yang *et al*., 2021). In both pear and grapevine buds, chilling-mediated dormancy release has been linked to a decline in ABA levels, which corresponds with downregulation of ABA biosynthetic genes and upregulation of in ABA catabolism genes (Zheng *et al*., 2015; Tuan *et al*., 2017). Exogenous applications of various plant hormones or their analogs has been shown to disrupt the natural deacclimation process in plants (Jacobsen *et al*., 2013; Zheng *et al*., 2018b; Kovaleski & Londo, 2019; Yuxi *et al*., 2021; Wang *et al*., 2022, 2025). A group of MADS-box transcription factors, known as DORMANCY-ASSOCIATED MADS-box genes (DAMs), have been noted for their potential regulation of dormancy transition under chilling accumulation. In *Prunus persica*, six tandemly arranged DAM genes located at the end of chromosome 1 were identified, and a deletion at this locus was associated with the ‘evergrowing’ phenotype (Bielenberg *et al*., 2008). The expression of all six copies of DAM is highly correlated with chilling accumulation (Li *et al*., 2009; Yamane *et al*., 2011; Yu *et al*., 2020). In addition, chilling during the dormant season was also found to induce DNA methylation changes, particularly in the CHH context, in perennial crops like apple (*Malus domestica*) and sweet cherry (*Prunus avium*), regulating gene expression for dormancy release and growth preparation (Kumar *et al*., 2016; Rothkegel *et al*., 2020). While several physiological studies have explored the relationship between chilling accumulation and dormancy progression, the underlying molecular mechanisms remain poorly understood.

In this study, we conducted an experiment involving five field collections of grapevine buds across the dormant season, each representing a different level of accumulated chilling. Following each collection, buds were subjected to a controlled forcing assay simulating growth-permissive conditions to capture dynamic transcriptomic responses to warm temperature exposure and previous field chilling. Our objectives were to: (1) investigate the molecular basis of the dormant bud response to warm temperature; (2) determine whether these responses differ according to previous field chilling; and (3) identify the portion of the transcriptome that is responsive to previous field chilling only, independent of warming. We hypothesized that chilling alters the hormone signaling and transcriptional readiness in a way that reactivates temperature sensitivity in dormant buds, enabling deacclimation associated growth-related pathways upon exposure to warm conditions. This research provides critical insights into the molecular transition from endodormancy to ecodormancy and is essential for improving climate resilience in perennial crop systems.

## Results

### Grapevine cold hardiness dynamics under field condition

We conducted a time-series experiment to assess the physiological and transcriptomic responses of *Vitis vinifera* ‘Cabernet Sauvignon’ grapevines collected from the field during the 2021-2022 dormant season. Cane samples were collected on November 3 (Collection 1), November 16 (Collection 2), December 1 (Collection 3), December 14, 2021 (Collection 4), and February 20, 2022 (Collection 5) (Figure 1A). The ambient temperatures at the time of collection were 5.2°C, 2.7°C, 1.8°C, 2.1°C, and −4.1°C, respectively (Figure 1B). Field cold hardiness of the grapevine buds at each field collection date was measured as −11.4°C, −15.5°C, −18.8°C, −19.8°C, and −21.9°C from Collections 1 through 5, respectively (Figure 1B). To estimate chilling accumulation in the field, we applied the (Utah) chilling unit model (Shaltout & Unrath, 1983), which resulted in 394, 557, 677, 796, and 1044 chilling units for Collections 1 through 5, respectively (Figure 1C).

**Figure 1.**
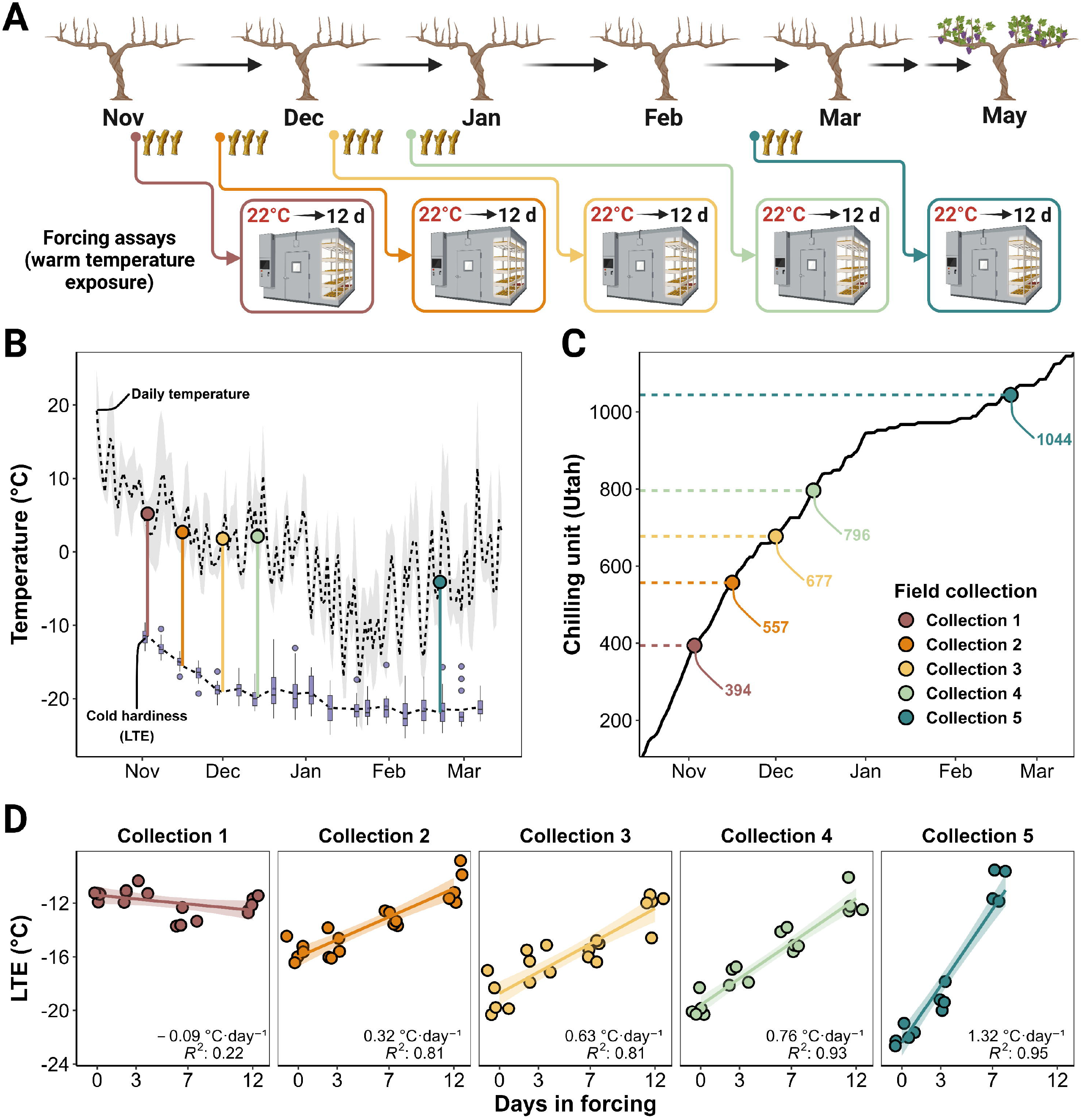
Experimental design, sample collection, data collection, and deacclimation assays. A) Schematic overview of the experimental design and sampling timeline; B) Daily ambient temperature throughout the 2021–2022 dormant season, along with corresponding field temperature and bud cold hardiness at each sampling timepoint; C) Chilling unit accumulation based on the NC model and corresponding chilling units at each sampling timepoint; D) Bud cold hardiness in the forcing assays and the computed *k*_*deacc*_.

### Diverged deacclimation under growth-permissive condition

To examine deacclimation response (*k*_*deacc*_), cane samples collected from the field were chopped into single-bud cuttings placed with cut ends in cups of DI water and incubated under growth-permissive forcing conditions. All sample collections exhibited significant deacclimation responses (Figure 1D) except those from Collection 1. Instead, a negative *k*_*deacc*_ slope was observed for samples in Collection 1, and the regression between cold hardiness and time in forcing showed minimal fit (lowest R^2^; Figure 1D). Samples from Collections 2 through 5 showed increasing deacclimation capacity, with all corresponding regressions showing R^2^ values above 0.8. The calculated *k*_*deacc*_ steadily increased with chilling accumulation, ranging from 0.42 □ °C·d^−1^ in Collection 2 to 1.42□ °C·d^−1^ in Collection 5.

### Transcriptomic characterization of bud samples during controlled deacclimation

In parallel with each deacclimation assay, replicate buds were analyzed using RNA-seq to characterize their transcriptomic response to forcing temperatures and varying levels of chilling accumulation. A total of 95 RNA-seq libraries were generated from the five deacclimation assays. All libraries passed FastQC per-base quality standards, with an average of 5.2 million reads per library and a mean unique mapping rate of 81.2%. One library was identified as an outlier based on principal component analysis (PCA) and was excluded from downstream analyses (Figure S1). After filtering out low count genes (defined as fewer than ten transcripts per library), the final dataset consisted of 94 libraries and 10,428 genes, 24.6% of the total annotated grapevine transcriptome.

To characterize transcriptomic trends, PCA and weighted gene co-expression network analysis (WGCNA) were applied to the filtered expression matrix. PC1 and PC2, accounted for 27.2% and 11.2% of the total variance, respectively, differentiating libraries based on field chilling history and response to forcing conditions (Figure 2A). Transcriptomic profiles within each field collection were largely similar when examining timepoints collected in the first 4 h of exposure to forcing conditions (0 h, 2h and 4h). In contrast, by 24 h, libraries from all field collections shifted substantially toward the upper left quadrant of the PC1/PC2 plot, suggesting a transcriptomic transition between the within a day (0 to 4 h) and after a day (≥24 h) sample points, regardless of initial field conditions (Figure 2A). Libraries from Collections 1 to 4 exhibited limited transcriptomic change after 24 h (minimal separation between 24 h, 3 d, and 7 d libraries) (Figure 2A). In contrast, Collection 5 libraries continued to shift progressively toward the lower right region of the PCA plot after 24 h (Figure 2A).

**Figure 2.**
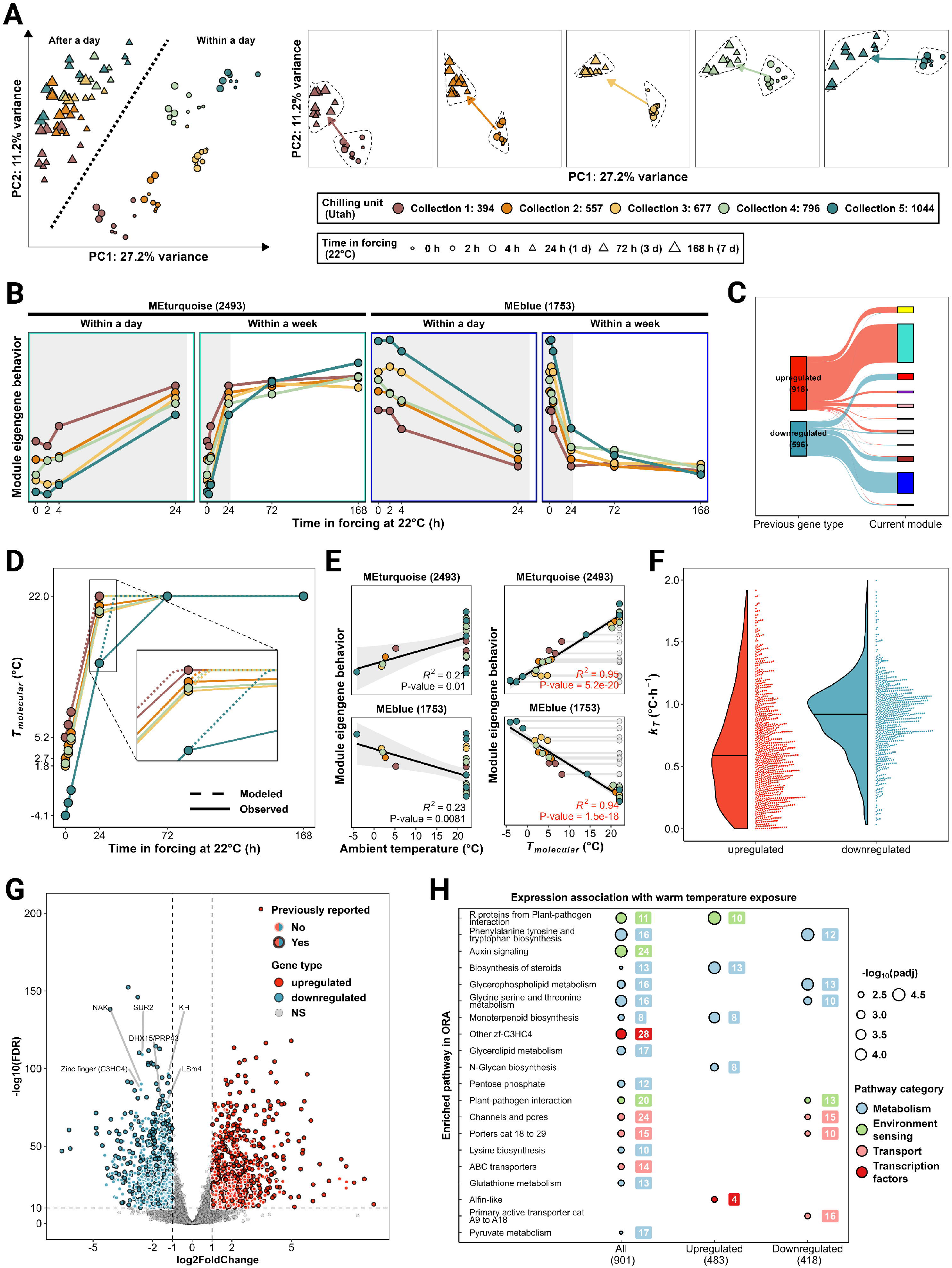
Transcriptomic response to warm temperature exposure. A) PCA of the active transcriptome, showing PC1 and PC2; B) Expression patterns of the MEs of co-expression modules ‘turquoise’ and ‘blue’, which only responded to warm temperature exposure.; C) Distribution of previously identified temperature-responsive genes across WGCNA-defined modules; D) Computed *T*_*molecular*_ of each sample, derived from the *k*_*T*_ of module ‘turquoise’; E) Correlation between MEs of modules ‘turquoise’ and ‘blue’ with both ambient temperature and *T*_*molecular*_; F) *k*_*T*_ of individual temperature-responsive genes; G) Volcano plot of differentially expressed temperature-responsive genes; H) Pathway enrichment analysis of the identified temperature-responsive genes.

The WGCNA using all the genes in the final gene count matrix identified ten gene co-expression modules, excluding module ‘grey’ containing noisy genes. The number of genes in each module ranged from 307 in module ‘purple’ to 2493 in module ‘turquoise’. The module eigengenes (MEs) of all the gene co-expression modules are depicted in Figure S2.

### Transcriptomic response to warm temperature exposure

The initial RNA-seq analysis focused on the response to the forcing conditions and assessing whether this response differed depending on field chilling accumulation. The MEs of the two largest gene co-expression modules ‘turquoise’ and ‘blue’ exhibited distinct dynamic behaviors during the forcing assays. These trends were closely mirrored or inversely correlated with PC1 from PCA (Figure 2B and Figure S3). Together, these two modules represented nearly 40% of the active transcriptome in our study. Although the initial ME expression levels under field conditions differed across collections, a common temporal response pattern emerged across all field collections. Little change was observed within the first 2 h of forcing, but a slight increase/decrease occurred by 4 h. The greatest transcriptomic shift occurred between 4 h and 24 h, and the overall magnitude of change during this period was consistent across all collections, regardless of previous field conditions. After 24 h, samples from Collections 1 through 4 showed minimal additional change in ME expression at 3 d and 7 d. In contrast, the ME expression in Collection 5 samples increased/decreased between 24 h and 3 d (Figure 2B).

Temperature-responsive genes identified in a previous study (Wang *et al*., 2024b) were compared with the genes associated with the WGCNA co-expression modules from this study. Among the 918 genes previously identified as positively correlated with temperature, 660 (72%) were assigned to the module ‘turquoise’ (Figure 2C). Among the 596 genes previously identified as negatively correlated with temperature, 357 (60%) were assigned to module ‘blue’. These suggest that modules ‘turquoise’ and ‘blue’ collectively captured temperature-responsive genes in our dataset.

### Molecular temperature model

Based on this observation, we developed an empirical gene expression model to describe transcriptomic adaptation to warm temperature exposure during the forcing assays. The model assumes that when ambient temperature increases to 22°C, gene expression shifts linearly toward a target expression level with a constant molecular temperature response rate (*k*_*T*_), independent of previous field chilling. Using ME expressions from the ‘turquoise’ and ‘blue’ modules, we estimated *k*_*T*_ through the empirical gene expression model (Equation 1). For the module ‘turquoise’ that represents a positive shift in gene expression towards the 22°C target, the *k*_*T*_was 0.76□°C·h□^1^. For the module ‘blue’ that represents a negative shift in expression, the *k*_*T*_ was 0.77□°C·h□^1^. These rates were subsequently used to calculate the molecular temperature (*T*_*molecular*_), which theoretically aligns with current expression of the temperature-responsive genes, as a function of baseline temperature under field condition, time in forcing, and the target temperature of 22°C (Equation 2 and Equation 3). As calculated, the *T*_*molecular*_ plateaued at approximately 23 h for Collection 1, while Collections 2 through 4 reached their target expression levels shortly after 24 h (Figure 2D). In Collection 5, the *T*_*molecular*_ continued to change and reached stability around 35 h (Figure 2D). To validate this model, we performed correlation analysis comparing ME expression values of modules ‘turquoise’ and ‘blue’ to both ambient temperature and *T*_*molecular*_. Regression using the *T*_*molecular*_ yielded significantly improved fit (higher R^2^ and lower p-values; Figure 2E).

Using *T*_*molecular*_ as the only numeric factor in differential expression analysis, we identified 1,820 differentially expressed genes (DEGs) with highly significant expression correlations (padj ≤ 1e-10; Data S1). Of these, 1,029 were upregulated and 791 were downregulated in response to warm temperature exposure. The *k*_*T*_ of individual upregulated genes ranged from 0 to 2□°C·h□^1^, with an average of 0.66□°C·h□^1^, and a distribution skewed toward 0 to 0.5□°C·h□^1^. To compare, the *k*_*T*_ of individual downregulated genes followed a normal distribution within the same range, with a higher average of 0.96□°C·h□^1^ (Figure 2F).

Many of the most significantly differentially expressed genes with the greatest log□ fold changes overlapped with the previously identified temperature-responsive genes (Wang *et al*., 2024b) (Figure 2G). However, among the downregulated genes, we identified several novel and highly significant members not previously reported. These include *VIT_01s0026g02310* (encoding Sphinganine C4-monooxygenase/SUR2), *VIT_14s0108g01040* (KH domain-containing protein), *VIT_02s0012g00150* (Numb-associated kinase/NAK), *VIT_17s0000g06610* (Zinc finger protein C3HC4), *VIT_07s0005g01770* (DEAH-box RNA helicase/DHX15), and *VIT_07s0005g00230* (U6 snRNA-associated Sm-like protein/LSm4). Pathway enrichment analysis of the temperature-responsive genes revealed 20 significantly enriched pathways primarily related to the Metabolism and Environmental Sensing categories (Figure 2H).

### Transcriptomic response to chilling

The second analysis aimed at deciphering the impact of previous field chilling accumulation on the gene expression during the forcing assays. We identified several co-expression modules whose ME showed expression patterns that either correlated with previous field chilling or reflected the interactions of chilling and forcing exposure.

We identified one co-expression module ‘black’ that was negatively correlated with chilling accumulation but exhibited no interaction effects during deacclimation assays (Figure 3A). Pathway enrichment analysis of the identified 565 annotated DEGs that showed this behavior (out of a total of 1,138 DEGs; Data S2) revealed eight significantly enriched pathways (Figure 3B). In the protein processing in the endoplasmic reticulum pathway, 14 out of 16 DEGs negatively correlated with chilling and were associated with heat shock proteins (HSPs). Further examination of all the 1,138 chilling-responsive DEGs expanded the list to include 21 DEGs that are associated with HSPs or heat shock transcription factors (HSFs). The expression of these genes showed a significant and consistent negative correlation with previous chilling (Figure S4). Given the large number of DEGs associated with HSPs and HSFs, we examined the expression pattern for all genes encoding HSPs and HSFs in the current transcriptome annotation. Nearly two-thirds of all HSP genes followed the same expression behavior as the ME of module ‘black’ (Figure 3C).

**Figure 3.**
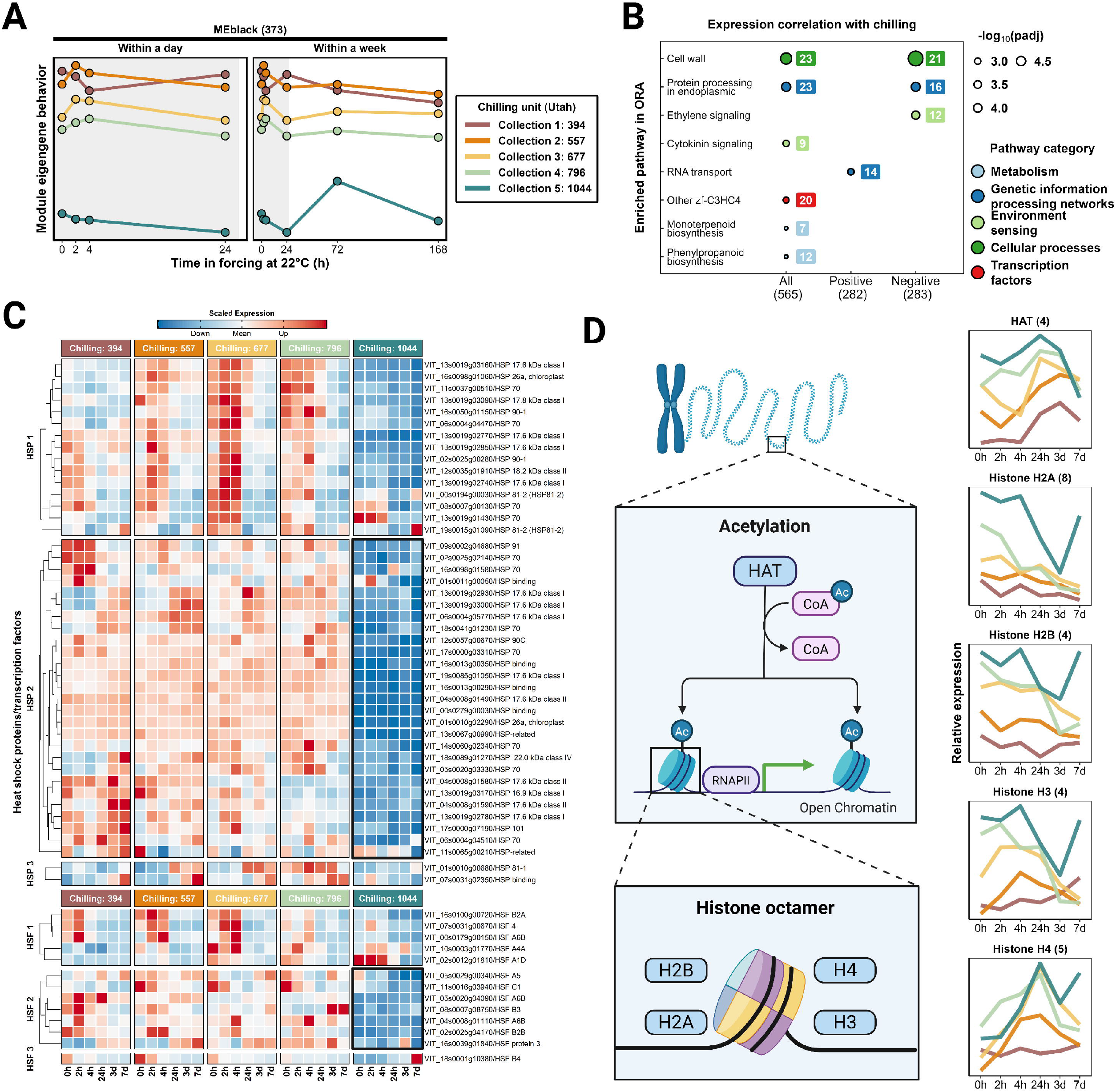
Transcriptomic response to chilling. A) Expression pattern of the ME of module ‘black’ that showed chilling only response; B) Pathway enrichment analysis of the chilling-responsive genes; C) Expression of all genes encoding HSPs and HSFs; D) Schematic of histone acetylation and the summed expression of genes encoding histone octamer components (H2A, H2B, H3, and H4) and HATs. Numbers beside protein names indicate the number of gene copies included in the expression sum.

A second enriched group of genes correlated with field chilling was histone-related proteins. These 13 DEGs encode H2A (2 DEGs), H2B (3 DEGs), H3 (3 DEGs) and H4 (5 DEGs) genes and their expression pattern was the inverse pattern of the ME module ‘black’ (Figure S5). Using the same approach, we investigated the expression patterns of all genes encoding histone-related proteins and found that the summed expression of all the genes encoding for the proteins in histone octamer: H2A (eight copies), H2B (four copies), H3 (four copies) and H4 (five copies) and histone acetyltransferase (HAT, four copies) also followed this pattern (Figure 3D).

### Transcriptomic response to the interaction of chilling and warm temperature exposure

Next, we evaluated the impact of the interaction of warm temperature exposure and previous field chilling accumulation on gene expression dynamics under forcing conditions. We observed that two co-expression modules, ‘brown’ and ‘yellow’, were correlated with chilling accumulation and subsequently shifted during the forcing assays. For module ‘brown’, ME expression was positively correlated with chilling under field conditions, remained stable during the first 24□h of forcing, and then progressively decreased in higher-chilling samples while slightly increasing in lower-chilling samples, resulting in convergence by 7 d. In contrast, module ‘yellow’ was negatively correlated with chilling under field conditions but exhibited the opposite trend during forcing: ME expression increased in high-chilling samples and decreased in low-chilling ones, also converging by 7 d. These dynamics reveal chilling-dependent patterns in response to warm temperature, which we describe as **chilling-positive convergence** (module ‘brown’) and **chilling-negative convergence** (module ‘yellow’), both characterized by initial chilling-associated divergence followed by convergence during the forcing assay (Figure 4A).

**Figure 4.**
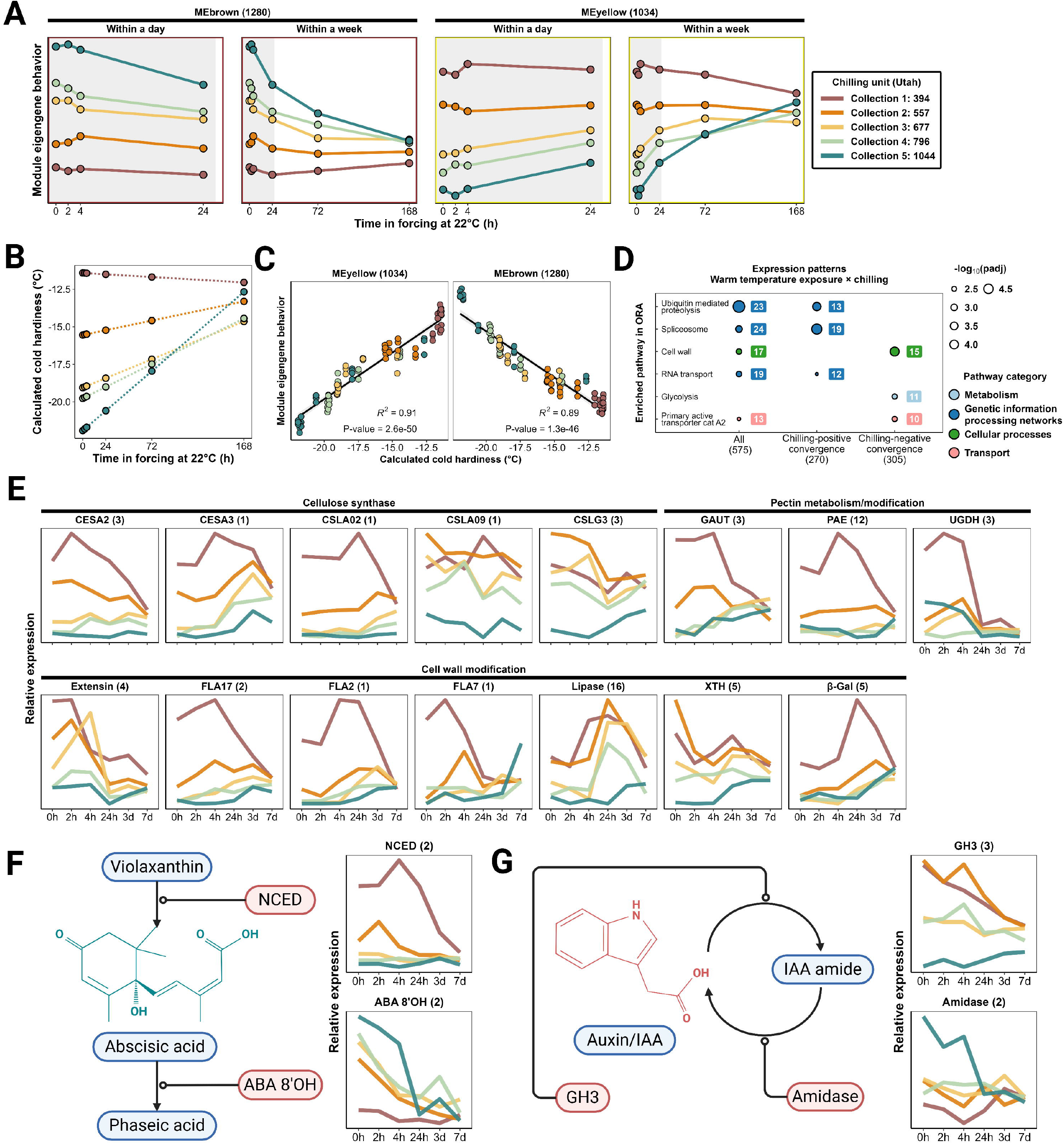
Transcriptomic response to the interaction of chilling and warm temperature exposure. A) Expression patterns of MEs for modules ‘brown’ and ‘yellow’, showing ‘chilling-positive convergence’ and ‘chilling-negative convergence’ patterns, respectively; B) Calculated cold hardiness of buds during the experiment; C) Correlation of MEs from modules ‘brown’ and ‘yellow’ with calculated cold hardiness; D) Pathway enrichment analysis of the chilling-positive convergence and chilling-negative convergence genes; E) Summed expression of genes encoding cell wall proteins that followed the chilling-negative convergence pattern; F) Schematic of abbreviated ABA metabolism and the summed expression of NCED and ABA 8’OH genes; G) Schematic of auxin-amino acid conjugation and deconjugation, and the summed expression of GH3 and amidase gene. Numbers beside protein names indicate the number of gene copies included in the expression sum.

We also observed that the behavior of these MEs (chilling-positive convergence and chilling-negative convergence) appeared to be closely associated with bud cold hardiness. Using the observed *k*_*deacc*_ and the initial cold hardiness values for each collection, we calculated cold hardiness at each timepoint during the five deacclimation assays (Equation 4, Figure 4B). The ME expression patterns of modules ‘brown’ and ‘yellow’ exhibited strong and statistically significant correlations with the calculated cold hardiness values across timepoints (Figure 4C).

Pathway enrichment analysis of the annotated DEGs that showed the ME patterns of module ‘brown’ and ‘yellow’ (Data S3) identified six overrepresented pathways (Figure 4D). Among the 17 DEGs associated with the overrepresented cell wall pathway, 15 exhibited chilling-negative convergence behavior. We then examined the overall expression patterns of all cell wall-related genes that passed low-count filtering. This analysis revealed that, among 33 annotated proteins in the cell wall pathway, 13 exhibited an expression pattern that matched either the chilling-negative convergence pattern or the chilling only response pattern. The total expression of the genes encoding these proteins was higher under field conditions in early collections (lower chilling), followed by a gradual convergence toward similar expression levels across all collections as the forcing assay progressed (Figure 4E).

In addition, although not statistically overrepresented in the pathway enrichment analysis, several genes involved in ABA and auxin metabolisms exhibited expression patterns associated with the interaction between warm temperature exposure and chilling. In the ABA metabolism pathway, the summed expression of 9-cis-epoxycarotenoid dioxygenase (NCED, key ABA biosynthesis enzyme) genes was highest in early collections under field conditions but declined and eventually converged to a minimum level across all collections by 7 d. Conversely, the summed expression of ABA 8’-hydroxylase (ABA 8’OH, key ABA degradation enzyme) genes was highest in later collections under field conditions and similarly decreased over time, converging by 7 d. (Figure 4F). In the auxin metabolism pathway, the summed expression of Gretchen Hagen 3 (GH3, key auxin conjugation enzyme) genes and amidase genes (key auxin de-conjugation enzyme) followed the chilling-negative convergence pattern and chilling-positive convergence pattern, respectively (Figure 4G).

### Plant hormone signaling pathways and ICE, CBF/DREB1 genes

We manually examined the expression of the genes involved in several plant hormone signaling pathways that have been previously reported to be correlated with grapevine cold hardiness and deacclimation (Chai *et al*., 2019; Londo & Kovaleski, 2019; Kovaleski & Londo, 2019; Wang *et al*., 2024b) and the members of the ICE and CBF/DREB1 genes (Xiao *et al*., 2006; Noriega *et al*., 2024).

In the ethylene signaling pathway, the overall expression of signaling components did not show a consistent response to chilling. However, several members, including Ethylene Insensitive 3 (EIN3), Mitogen-Activated Protein Kinase 3 (MPK3), Constitutive Triple Response 1 (CTR1) and EIN3-Binding F-box Protein 1 (EBF1) responded rapidly to warm temperature exposure under the forcing assays. We also identified six ethylene responsive transcription factors (AP2/ERFs) that exhibited a negative correlation with previous field chilling (Figure S6), which aligns with the overrepresentation of ethylene signaling among the DEGs that showed the chilling only response (Figure 3B). In the ABA signaling pathway, the overall expression of the genes encoding Protein Phosphatase 2C (PP2C, eight genes) showed a general chilling-only response, being negatively correlated with field chilling. The overall expression of the genes encoding SNF1-Related Protein Kinase 2 (SnRK2, four genes) showed a chilling-positive convergence pattern (Figure S7). In the auxin signaling pathway, the overall expression of multiple auxin-responsive transcription factors (ARFs) showed clear response to warm temperature exposure, though in both directions: some increased and others decreased under forcing conditions. Similarly, components of auxin receptors and signaling intermediates exhibited rapid but divergent responses to warm temperature exposure (Figure S8). Only three genes in the ICE and CBF/DREB1 gene group passed the low count filtering. The expression of the two DREB genes, *VIT_11s0016g02140* and *VIT_13s0067g01960*, displayed a general chilling-positive convergence behavior (Figure S9).

## Discussion

Warming and increasingly unstable winters driven by climate change may threaten the sustainability of global grape and wine production by disrupting the physiological progression of grapevine dormancy. Although the physiological and transcriptomic mechanisms underlying cold acclimation and deacclimation have been a topic of recent studies (Chai *et al*., 2019; Kovaleski & Londo, 2019; Camargo-Alvarez *et al*., 2020; North & Kovaleski, 2023; Wang *et al*., 2024b,c; Noriega *et al*., 2024; Londo & Kovaleski, 2025; De Rosa *et al*., 2025; Wang *et al*., 2025; Guo *et al*., 2025), far less is known about how field chilling accumulation shapes transcriptomic responsiveness during the dormant season. This study aimed to uncover how changing chilling accumulation influences the gene expression responses of buds during deacclimation in forcing conditions.

In this study, ‘Cabernet Sauvignon’ grapevine buds were sampled from the field five times (Collection 1 to 5) during the dormant season. The chilling accumulation of the five collections ranged between 394 and 1044 hours (Utah model). Based on the previously established relationship between chilling accumulation and deacclimation potential in ‘Cabernet Sauvignon’ (Londo & Kovaleski, 2025), buds were estimated to have reached approximately 0%, 12%, 25%, 40%, and 75% of their maximum deacclimation potential at Collections 1 through 5, respectively. Consistent with these estimates, the *k*_*deacc*_ increased proportionally during the deacclimation assays, further supporting the chilling-mediated progression of deacclimation potential (Londo & Kovaleski, 2025). Based on the chilling requirement of ‘Cabernet Sauvignon’, buds in Collection 1 were endodormant, in transition from endodormancy to ecodormancy in Collection 2 through 4, and ecodormant in Collection 5 (Londo & Johnson, 2014; Kovaleski *et al*., 2018; Londo & Kovaleski, 2025).

The gene expression analysis conducted here revealed new insights into the regulation of chilling, warm temperature sensing, and their interaction under growth-permissive conditions where deacclimation rate varies. The majority of genes from the four largest modules in the this study, ‘turquoise’, ‘blue’, ‘brown’, and ‘yellow’, were broadly classified as temperature-responsive in a previous study using weekly RNA-seq sampling (Wang *et al*., 2024b). However, the modules ‘turquoise’ and ‘blue’ clearly displayed a much faster transcriptomic response to the warm temperature exposure, with most changes occurring within the first 24 h across all five field collections, regardless of previous field chilling accumulation. In contrast, modules ‘brown’ and ‘yellow’ showed a slower mode of action, where both the rate and the direction of expression change were impacted by previous chilling, resulting in a delayed expression convergence by 7 d. This clear differentiation enables the separation of temperature-responsive genes (genes in modules ‘turquoise’ and ‘blue’) from those genes with responses modulated by chilling (genes in modules ‘brown’ and ‘yellow’). Although several studies have explored transcriptomic changes during grapevine deacclimation (Kovaleski & Londo, 2019; Wang *et al*., 2022; Noriega *et al*., 2024; De Rosa *et al*., 2025), this is the first study to incorporate field chilling accumulation as a factor in the analysis. This approach enabled the identification of genes that respond specifically to chilling, independent of warm temperature exposure, such as those in module ‘black.

To further explore the expression pattern of the temperature-responsive gene, we conducted empirical modeling and differential expression analysis based on computed *T*_*molecular*_ for genes that respond rapidly to warm temperature exposure. Analysis of module eigengene (ME) behavior revealed three key insights: (1) minimal transcriptomic changes occurred within the first 4 h; (2) most gene expression changes occurred between 4 and 24 h after warming; and (3) the magnitude and rate of transcriptomic change were largely independent of field sampling time, indicating a similar response to warming across different previous field chilling accumulation. Notably, the maximum temperature increase in this experiment was 26°C (from the coldest field sample to the growth chamber condition), and most of the gene expression changes were completed within one day. This suggests that in dormant grapevine buds, simple temperature-responsive gene regulation is rapid and chilling-independent.

Based on this behavior, we calculated a theoretical molecular temperature response rate, termed *k*_*T*_, as 0.76□°C·h□^1^ for upregulated and 0.77□°C·h□^1^ for downregulated gene modules, respectively. These rates suggest a maximum transcriptomic adjustment capacity of approximately 18°C per day. Therefore, under natural conditions, when ambient temperatures fluctuate more than 18°C within a single day, the transcriptomic machinery in dormant grapevine buds may be unable to fully adjust. Since transcriptional regulation often represents the earliest molecular response to environmental stimuli (Nievola *et al*., 2017), such mismatches between diurnal temperature shifts and transcriptomic responsiveness may lead to insufficient physiological responses, reducing the plant’s ability to adjust to environmental stress in a timely manner. As rapid and extreme temperature fluctuations are projected to become more frequent and intense under future climate scenarios (Wu *et al*., 2025), further research is urgently needed to evaluate how large winter temperature swings disrupt the physiological processes in the dormant season.

Using the empirically determined *k*_*T*_, we also calculated the *T*_*molecular*_ for each RNA-seq library, which served as a proxy variable for identifying temperature-responsive genes. A total of 1,820 DEGs exhibited highly significant correlations with *T*_*molecular*_, including 1,029 upregulated and 791 downregulated in response to warm temperature exposure. The abundance of highly significant temperature-responsive genes identified in this study indicates that a substantial portion of the active transcriptome of grapevine bud in the dormant season is associated to temperature sensing and response, and that this regulatory mechanism remains active regardless of the dormancy status. While the overall behavior aligned with the eigengene patterns of modules ‘turquoise’ and ‘blue’, we observed a slight difference in *k*_*T*_ at the individual gene level. Notably, genes that were downregulated in response to warm temperature exposure, on average, a 31% higher *k*_*T*_ compared to the upregulated genes. If this relationship holds true under natural conditions, it suggests that during a cold event, cold-induced genes are upregulated more rapidly than cold-suppressed genes are downregulated. This asymmetric response may reflect underlying biological mechanisms, where the suppression of gene expression under cold may be partially passive (e.g., reduced enzymatic activity), whereas the induction of cold-responsive genes involves a faster active transcriptional programming (Ding *et al*., 2024).

Most of the temperature-responsive genes identified in this study have been reported in a previous study (Wang *et al*., 2024b), but we also identified six very significant new members that were downregulated under warm temperature exposure. With the exception of *VIT_02s0012g00150* (NAK gene), whose function in plants remains poorly characterized, the other five genes have been implicated in key biological processes in other plant species.

*VIT_01s0026g02310* encodes SUR2, a key enzyme in sphingolipid biosynthesis, a pathway implicated in cold stress signaling and membrane stabilization (Ali *et al*., 2018).

*VIT_14s0108g01040*, encodes a KH domain-containing RNA-binding protein, which has been associated with thermotolerance and heat-induced gene regulation through modulation of stress-responsive transcription factors (Guan *et al*., 2013). *VIT_07s0005g01770* encodes DHX15, and *VIT_07s0005g00230* encodes LSm4. These proteins are involved in RNA modification, and their downregulation may mark reduced demand for translational regulation under warming (Bouveret *et al*., 2000; Mosallanejad *et al*., 2014; Kerbler & Wigge, 2023). *VIT_17s0000g06610*, encodes a C3HC4-type RING zinc finger protein/E3 ligase and is likely involved in protein ubiquitination and degradation during abiotic stress. E3 ligases have been shown to regulate abiotic stress adaptation in different crops (Agarwal & Khurana, 2020; Han *et al*., 2021). Detailed functional characterization of these genes in grapevine is still needed, and their significant downregulation under warm temperature exposure suggests that they may play conserved roles in maintaining transcriptomic stability and dormancy-associated signaling.

Pathway enrichment analysis revealed that temperature-responsive genes are predominantly associated with metabolism and environmental sensing pathways as seen in a previous study (Wang *et al*., 2024b). These two pathway groups likely act synergistically: temperature sensing initiates metabolic reprogramming, while the resulting metabolic shifts can, in turn, modulate and reinforce stress signaling. Among all plant hormones, auxin was the only signaling pathway significantly overrepresented among the temperature-responsive genes. Detailed investigation revealed that eight ARFs, two auxin receptors, four negative regulators, and four positive regulators of auxin signaling exhibited expression patterns that were highly correlated with the *T*_*molecular*_. Although the direction of regulation varied (some were upregulated while others were downregulated under warm temperature exposure), this overall pattern suggests a critical role for auxin signaling in immediate temperature sensing and response, being consistent with previous findings (Gray *et al*., 1998; Bellstaedt *et al*., 2019). The rapid modulation of auxin signaling likely facilitates swift growth adjustments, such as thermomorphogenesis, and integrates with other regulatory networks to coordinate adaptation and the initiation of developmental transitions under changing thermal conditions (Bianchimano *et al*., 2023).

Another key insight from this study is the identification of three gene co-expression modules, ‘black’, ‘brown’, and ‘yellow’, that showed significant associations with chilling accumulation. The module associated with responses correlated with chilling alone (module ‘black’) included the cell wall pathway, protein processing in the endoplasmic reticulum, and ethylene signaling. The cell wall pathway was also significantly enriched among genes responsive to the interaction between warm temperature exposure and chilling (modules ‘brown’ and ‘yellow’) with nearly 40% of the pathway’s proteins followed either a chilling-only (negative) pattern or a chilling-negative convergence pattern. Cell wall remodeling has long been recognized as a key mechanism in chilling-mediated dormancy progression. For example, in apple (*Malus domestica*), DNA methylation regulates genes like *Pectinesterase 3*, which promotes pectin remodeling critical for cell wall flexibility during chilling-induced dormancy release (Kumar *et al*., 2016). In *Polygonatum kingianum*, chilling has been shown to cause structural changes in plasmodesmata and the cell wall, supported by transcriptomic evidence for the upregulation of cell wall-loosening genes (Wang *et al*., 2019). Similarly, in *Arabidopsis thaliana*, chilling and sub-zero acclimation trigger broad alterations in cell wall composition, including the induction of enzymes such as pectin methylesterases and xyloglucan endotransglucosylase/hydrolases like XTH19, which are important for structural remodeling (Takahashi *et al*., 2019, 2021). In grapevines, the upregulation of another group of genes encoding cell wall proteins were found intensively upregulated with other growth-related pathways when approaching budbreak (Wang *et al*., 2025). These changes are believed to enhance intercellular communication and prime tissues for growth reactivation post-dormancy (Alonso Baez & Bacete, 2023). Our findings suggest that intensive cell wall remodeling occurs during the early dormant season in grapevine buds, and this remodeling diminishes as chilling accumulates. Warm temperature exposure further reduces the intensity of this process. This observation suggests that unseasonal warming events in early winter may suppress cell wall remodeling activity, potentially disrupting normal dormancy cycling (Kovaleski, 2024).

Genes encoding HSPs and heat HSFs exhibited expression patterns negatively associated with chilling yet remained unresponsive to warm temperature exposure. Notably, the majority of these HSPs were small HSPs, which function as molecular chaperones, stabilizing and refolding proteins during stress conditions (Bourgine & Guihur, 2021). Although these proteins have been repeatedly reported for their expression correlation and stabilizing functionality under cold stress in various plant species (Takemura & Tamura, 2016; ul Haq *et al*., 2019; Chang *et al*., 2021; Sadura & Janeczko, 2024), limited information is available for their potential function in the chilling-mediated dormancy progression. The genes encoding several histone-related proteins, including H2A, H2B, H3, H4 and HATs, were also found to be positively correlated with chilling. The upregulation of the gene encoding these proteins during the dormant season suggests an active chromatin remodeling process in grapevine buds. Similar histone modification has been found in poplar (Conde *et al*., 2019), aspen (Karlberg *et al*., 2010), chestnut (Santamaría *et al*., 2009), apple (Kumar *et al*., 2016) and grapevine (De Rosa *et al*., 2024), and has been proposed to be a key epigenetic mechanism underlying the chilling-mediated dormancy transitions by progressively enabling the onset of growth-related genes.

We found that the two modules that revealed the interaction between warm temperature exposure and chilling, ‘brown’ and ‘yellow’, were highly correlated with cold hardiness. This result suggests that a subset of the active transcriptome in dormant season grapevine buds integrates both chilling and immediate thermal cues to reflect their physiological state. In ABA metabolism, NCED (biosynthesis) genes decreased and ABA 8’OH (degradation) genes increased with chilling, suggesting a decline in ABA content, consistent with previous findings (Zheng *et al*., 2015, 2018a). Under forcing conditions, expression of both enzyme types dropped to minimal levels regardless of chilling history. Similarly, GH3 (auxin conjugation) decreased while amidase (deconjugation) increased with chilling, indicating a rise in free auxin, followed by convergence under forcing. Several ERF genes in ethylene signaling pathway also showed this pattern. Our analysis indicates, first, that chilling accumulation drives transcriptomic shifts across multiple hormone pathways, potentially reflecting a gradual, hormone-mediated dormancy progression.

Second, warm temperature exposure appears to override this chilling-dependent transcriptional regulation, which potentially resets the hormone-associated gene expression to a uniform level and turns transcriptomic switch from dormancy maintenance to growth reactivation.

Together, our findings provide a comprehensive examination of the transcriptomic dynamics underlying grapevine bud dormancy transitions and deacclimation. Based on the expression patterns observed in this study and existing knowledge of dormant season physiology, we propose a theoretical model to illustrate how chilling accumulation and warm temperature exposure regulate transcriptomic behavior in dormant buds (Figure 5). A large portion of the transcriptome is rapidly responsive to warm temperatures in a chilling-independent manner, which potentially indicates a conserved and constrained temperature sensing mechanism throughout the dormant season. These genes, largely involved in pathways of metabolism, environmental sensing, and auxin signaling, potentially form the core of the immediate temperature sensing and response mechanism. In contrast, chilling modulates the dormant-season transcriptome in two distinct ways. First, chilling-only responsive genes, such as those involved in histone modification and HSP synthesis, continuously change the internal state of bud under field conditions to facilitate dormancy progression. Second, the genes responsive to the interaction of chilling and warm temperature exposure, particularly those involved in ABA and auxin metabolisms and cell wall remodeling, could have a dual role in both dormancy progression in the field and chilling-dependent deacclimation/growth under growth-permissive condition.

**Figure 5.**
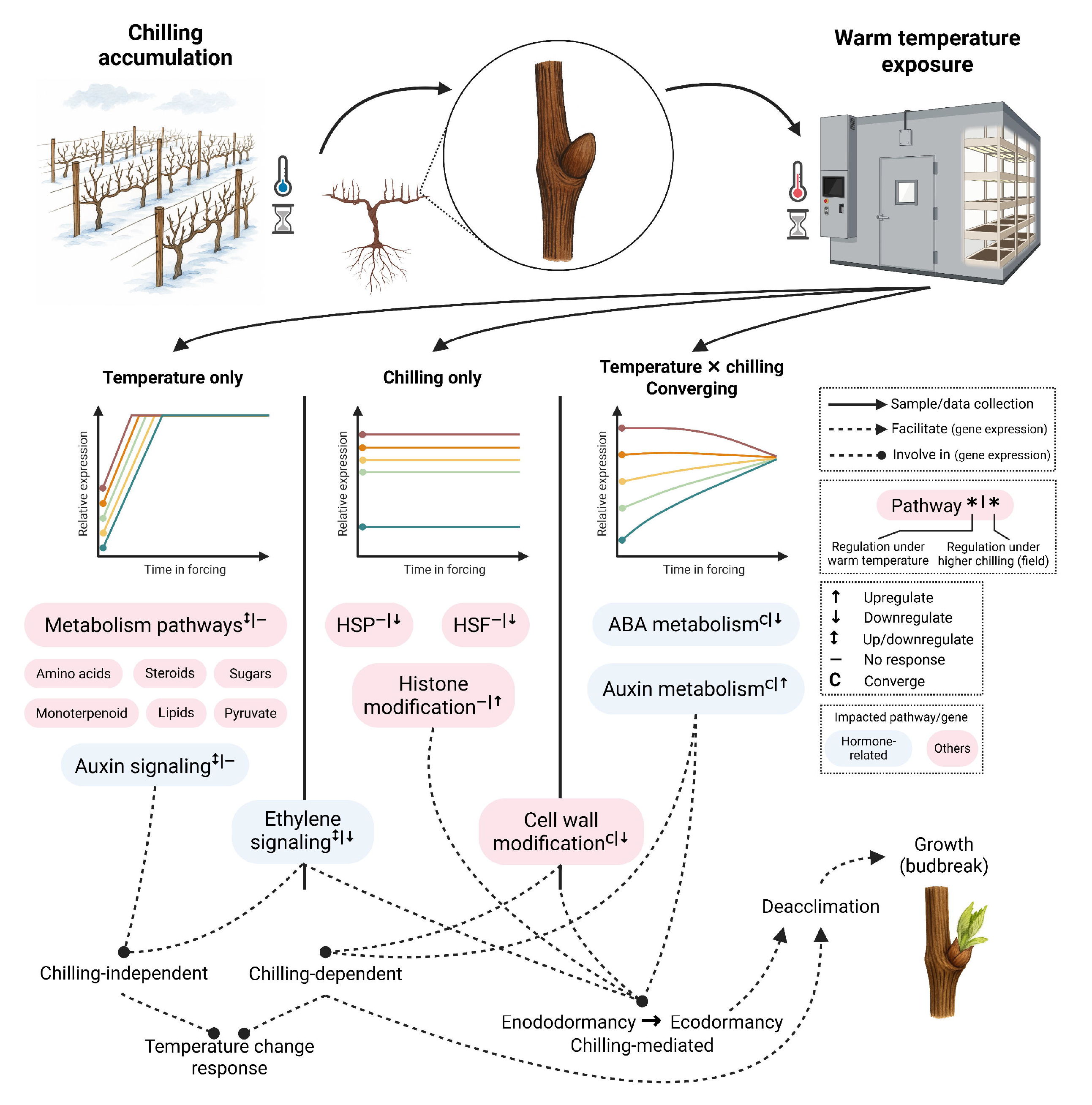
Conceptual model of chilling and warm temperature interaction in grapevine bud transcriptomic regulation. Schematic representation summarizing the roles of chilling accumulation and warm temperature exposure in regulating transcriptome during dormancy progression, deacclimation, and growth reactivation in grapevine buds.

Overall, the temperature-dependent, chilling-dependent, and chilling–temperature interaction gene expression behaviors observed in this study indicate the complex regulatory framework underlying grapevine dormant season physiology. By disentangling the effects of chilling and warm temperature exposure, we were able to uncover the transcriptomic basis of temperature sensing and the chilling-mediated dormancy transitions in grapevine. Several important questions remain. First, activation of the temperature-responsive genes identified in this study alone do not appear sufficient to induce physiological changes in cold hardiness (Wang *et al*., 2024b). How these genes contribute to preparing the plant for temperature adaptation, either independent or dependent of dormancy transitions, remains an open question for future research. Second, our experimental design assumes that chilling is the driver of dormancy transition.

However, because chilling accumulation is inherently confounded with time, and given ongoing debates about the accuracy of chilling models (including the Utah model used in this study) (Lin *et al*., 2022; North *et al*., 2024; Wang *et al*., 2024a), we cannot exclude the possibility that the observed gene expression changes are driven by chronological time rather than chilling per se. Future studies involving large-scale, high-resolution field RNA-seq datasets will be critical to distinguish chilling effects from temporal effects and to validate the concept of chilling-responsive transcriptomic regulation. Third, a substantial proportion of the genes showing significant correlation with chilling are unannotated, limiting our ability to interpret their functional relevance. Comprehensive annotation of these genes, potentially through comparative genomics and functional characterization, will be essential for fully uncovering the biology involved. Lastly, multi-omics validation (e.g., proteomics, metabolomics, and epigenomics) will be required to confirm the roles of candidate genes and to validate the findings from this study, which will help build a more mechanistic understanding of how temperature changes and chilling accumulation shape dormancy regulation in grapevines.

## Materials and Methods

### Plant materials and field sampling

Field-grown *Vitis vinifera* ‘Cabernet Sauvignon’ grapevines were used in this study. The experimental vines were commercially cultivated at Ravines Wine Cellars in Geneva, NY (42.845°N, 77.004°W). Vines were grafted onto 3309C rootstock and maintained under standard vineyard management practices. The experimental design and sample collection scheme are outlined in Figure 1. Dormant grapevine canes were collected at five timepoints during the 2021-2022 dormant season, November 3 (Collection 1), November 16 (Collection 2), December 1 (Collection 3), December 14, 2021 (Collection 4), and February 20, 2022 (Collection 5) to represent approximately equal chilling intervals spanning the transition from endodormancy to ecodormancy. Although there is a larger gap in calendar time between Collections 4 and 5, this period reflects the slower chill accumulation during mid-winter (Figure 1).

### Deacclimation forcing assays

After each field collection, canes were chopped into single-bud cuttings and incubated without supplemental light with cut ends in water and held at 22°C in a growth chamber. Such growth permissive forcing assay has been used to determine the deacclimation potential and the dormancy status of grapevine (Londo & Johnson, 2014; Wang *et al*., 2024b; Londo & Kovaleski, 2025). Bud cold hardiness was assessed at 0 h (field condition), 2 h, 4 h, 24 h (1 d), 72 h (3 d), and 168 h (7 d) after the start of forcing if the buds remained unbroken. Deacclimation rate (*k*_*deacc*_, in °C·h□^1^ or °C·d□^1^), the coefficient of the linear regression between bud cold hardiness and time under forcing conditions, was determined for each deacclimation assay conducted on the samples from each field collection (Kovaleski *et al*., 2018).

### Cold hardiness measurement

Bud cold hardiness was measured using differential thermal analysis (DTA) following standard protocols (Mills *et al*., 2006; Londo *et al*., 2023). Buds were excised along with surrounding cane tissue from single-node cuttings and placed into thermoelectric modules. Samples were then subjected to controlled freezing in a Tenney T2C environmental chamber (Tenney Environmental, PA, USA), programmed to cool at a rate of 4 °C·h^−1^ from 0°C to −50°C. The release of heat during tissue freezing, a phenomenon known as the low temperature exotherm (LTE) was detected and recorded via a Keithley 2700 data logger (Tektronix, Beaverton, OR, USA). The temperature at which LTE occurs corresponds to the cold hardiness of the grapevine bud (Pierquet & Stushnoff, 1980; Londo *et al*., 2023). Cold hardiness was measured for each timepoint of the deacclimation assays under growth chamber conditions, using five biological replicates per collection. In addition, field cold hardiness of ‘Cabernet Sauvignon’ grapevines from the same vineyard were continuously monitored throughout the dormant season to serve as a reference for evaluating deacclimation responses under controlled conditions.

### RNA-seq data collection

During each deacclimation assay, bud samples were collected in parallel with cold hardiness measurements for transcriptomic analysis using RNA-seq. For each timepoint, three biological replicates were prepared, each consisting of five pooled buds. Total RNA was extracted, and library construction was performed using the Lexogen QuantSeq 3’ mRNA-Seq Library Prep Kit (Lexogen, Greenland, NH), with technical support provided by the Cornell University Institute of Genomic Diversity (Ithaca, NY, USA). Sequencing was carried out at the Cornell University Institute of Biotechnology (Ithaca, NY, USA) using the Illumina NextSeq500 platform (Illumina, Inc., San Diego, CA, USA). Libraries were sequenced with a read length of 85 bp, and each library was sequenced in triplicate to ensure technical reproducibility.

### RNA-seq data analysis

RNA-seq data was analyzed following a previously generated pipeline specifically optimized for grapevine (Wang *et al*., 2022, 2024b). Standardized library QC (FastQC), trimming (BBDuk) was conducted for quality control of the libraries, and transcript alignment was conducted in STAR using *V. vinifera* 12X.v2 genome and VCost.v3 annotation (Dobin *et al*., 2013; Canaguier *et al*., 2017). DESeq2 was used to identify differentially expressed genes (Love *et al*., 2014).

PCA and WGCNA were used for the characterization of transcriptome and the identification of gene co-expression modules (Langfelder & Horvath, 2008). After the identification of target gene co-expression modules that revealed specific behavior under field or forcing conditions, we applied specific contrasting or correlation-based filtering to identify target genes.

To identify the genes that responded to the warm temperature exposure in the forcing assays (co-expression modules ‘turquoise’ and ‘blue’), we generated a molecular temperature model and calculated a molecular temperature response rate (*k*_*T*_) based on the eigengene expression patterns of the ‘turquoise’ and ‘blue’ co-expression modules. The calculation was performed as follows:

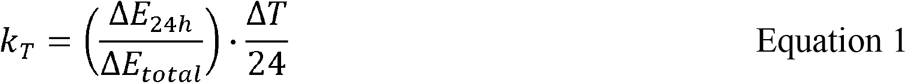

where *k*_*T*_ represents the molecular response rate to temperature change, expressed in °C·h^−1^; Δ*E*_24*h*_ is the change in eigengene expression in the Collection 5 samples during the first 24 h of forcing; Δ*E*_*total*_ is the total change in eigengene expression in the Collection 5 samples over the entire assay, calculated relative to the mean eigengene expression values on days 3 and 7 from all the samples; Δ*T* is the difference in temperature between the Collection 5 field condition and the constant 22°C growth chamber environment. The resulting *k*_*T*_ describes the rate at which gene expression responds to temperature change between the field condition and the controlled environment. The *k*_*T*_ was calculated separately modules ‘turquoise’ and ‘blue’, representing upregulated and downregulated genes under forcing conditions, respectively.

Assuming that temperature-responsive genes in grapevine respond linearly to changes in temperature, the molecular temperature that reveals the current expression at a given timepoint can be modeled using the molecular temperature response rate *k*_*T*_. The calculated molecular temperature *T*_*calc*_ is given by:

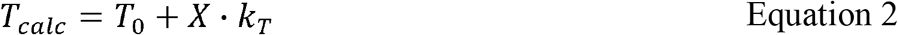

Where *T*_0_ is the baseline temperature under field condition, is the time under the forcing condition, *k*_*T*_ is the molecular response rate to temperature change. The actual molecular temperature at each timepoint, *T*_*molecular*_, is constrained by a temperature limit of 22°C in the forcing assay, and is defined as:

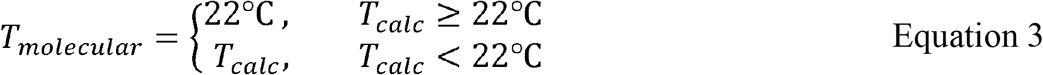

This formulation ensures that gene expression does not exceed its empirically observed target value while still capturing the linear response to temperature-driven deacclimation. Based on this formulation, the *T*_*molecular*_ was calculated for each field collection and growth chamber data collection timepoints and was used as the only numeric factor in a gene expression model in DESeq2 to identify these temperature responsive genes. Only the genes with a padj ≤ 1e-10 in such a model were identified as significant temperature responsive genes.

To identify genes that responded to previous field chilling (module ‘black’) or to the interaction between chilling and warm temperature exposure (modules ‘brown’ and ‘yellow’), we constructed a gene expression model using the formula: *design = ~ chilling + chilling × time_in_forcing*. For genes associated with both chilling and warm temperature exposure response (modules ‘brown’ and ‘yellow’), significance was determined by an padj ≤ 1e-5 for both chilling term and the interaction term. These genes were classified as significant temperature-responsive genes.

For genes that responded to chilling but did not change over time during the experiment (module ‘black’), significance was defined as padj ≤ 1e-5 for the chilling term and padj > 1e-3 for the interaction term, which specifies a chilling-specific but warm temperature-independent expression pattern.

In addition, based on the computed *k*_*deacc*_ and the measured initial cold hardiness, we also estimated the cold hardiness of the buds at each timepoint during the forcing assays. This calculation allowed us to assess the correlation between bud cold hardiness dynamics and the MEs of modules ‘brown’ and ‘yellow’ during deacclimation. The calculated cold hardiness (*CH*_*cal*_) at any timepoint is defined as:

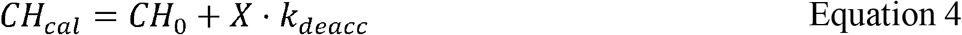

where *CH*_*cal*_ (°C) is the calculated cold hardiness, *CH*_0_ (°C) is the initial cold hardiness under field condition (0 h in the forcing assays), is the time under the forcing condition (in hours), *k*_*deacc*_ is the calculated deacclimation rate in °C·h□^1^ to facilitate the calculation of the cold hardiness within the first 4 h.

### Pathway enrichment analysis and other examinations

Enriched pathways among different identified target genes were detected through over representation analysis using VitisNet function pathways (Grimplet *et al*., 2009). Significantly enriched pathways at padj <= 0.005 were examined for correlations with known biological functions, and hub genes were determined based on the synergy of statistical significance and biological functionality. In addition, we also manually examined individual genes, or the genes involved in pathways that were previously reported to be functional in grapevine cold hardiness. To simplify interpretation and visualization, expression values of all gene copies encoding the same protein were summed in each figure, unless the discussion focused on individual genes.

Numbers in parentheses indicate the number of gene copies included in each sum.

## Supporting information

Supporting Material 1 will be used for the link to the file on the preprint site

Supporting Material 2 will be used for the link to the file on the preprint site

## Author contributions

JPL and HW designed and conducted the experiment. HW collected samples, conducted deacclimation assays, extracted RNA and conducted the RNA-seq analysis. JPL and HW wrote the manuscript.

## Acknowledgement

This research was funded in part by the 2022 Schmittau-Novak Grant from the School of Integrative Plant Science, Cornell University. We would like to acknowledge Hanna Martens and Felex Pike for helping with collecting dormant cuttings and in processing samples for cold hardiness analysis.

## Date availability

The RNA-seq raw data along with processed gene count matrix and sample metadata supporting the conclusions of this article are available in the NCBI-GEO repository accession GSE299692.

## Abbreviations

ABA: Abscisic acid
ICE-CBF-COR: Inducers of CBF Expression – C-Repeat Binding Factor/DRE Binding Factor – Cold Regulated Genes
*k*_*deacc*_: Deacclimation rates
DAM: DORMANCY-ASSOCIATED MADS-box gene
PCA: Principal component analysis
WGCNA: Weighted gene co-expression network analysis
PC: Principal component
ME: Module eigengene
*k*_*T*_: molecular temperature response rate
*T*_molecular_: molecular temperature
DEG: Differentially expressed gene
HSP: Heat shock protein
HSF: Heat shock transcription factor
HAT: Histone acetyltransferase
NCED: 9-cis-epoxycarotenoid dioxygenase
ABA 8’OH: ABA 8’-hydroxylase
GH3: Gretchen Hagen 3
EIN3: Ethylene Insensitive 3
MPK3: Mitogen-Activated Protein Kinase 3
CTR1: Constitutive Triple Response 1
EBF1: EIN3-Binding F-box Protein 1
AP2/ERF: Apetala 2/ethylene responsive transcription factor
PP2C: Protein Phosphatase 2C
SnRK2: SNF1-Related Protein Kinase 2
ARF: Auxin Response Factor
DTA: Differential thermal analysis
LTE: Low temperature exotherm

