## Supporting Material 1 will be used for the link to the file on the preprint site for "Time-series transcriptomics of grapevine deacclimation reveals chilling-dependent genetic responses to temperature increase during dormancy"

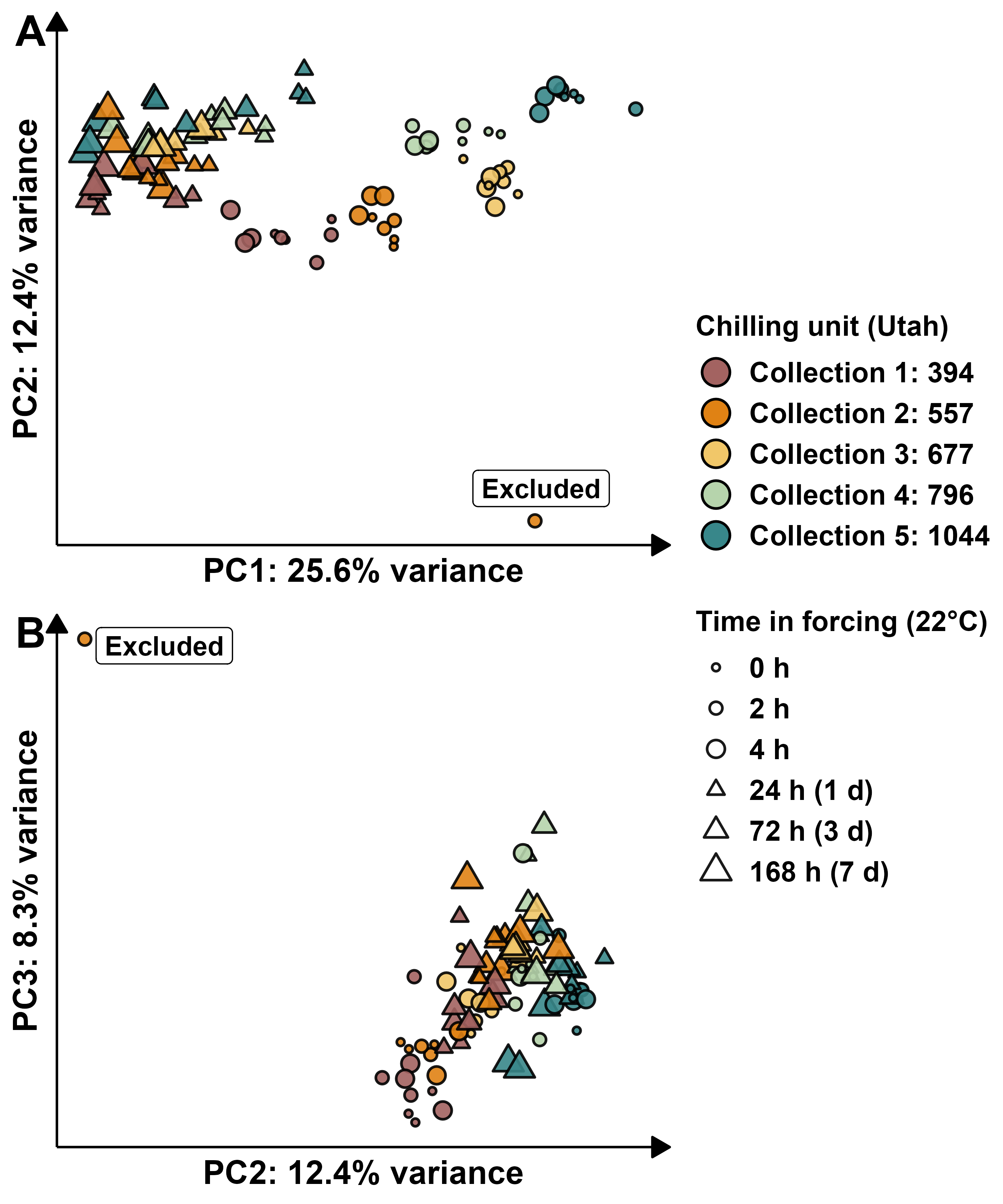

Figure S1. Identification of transcriptome outliers using Principal Component Analysis (PCA)

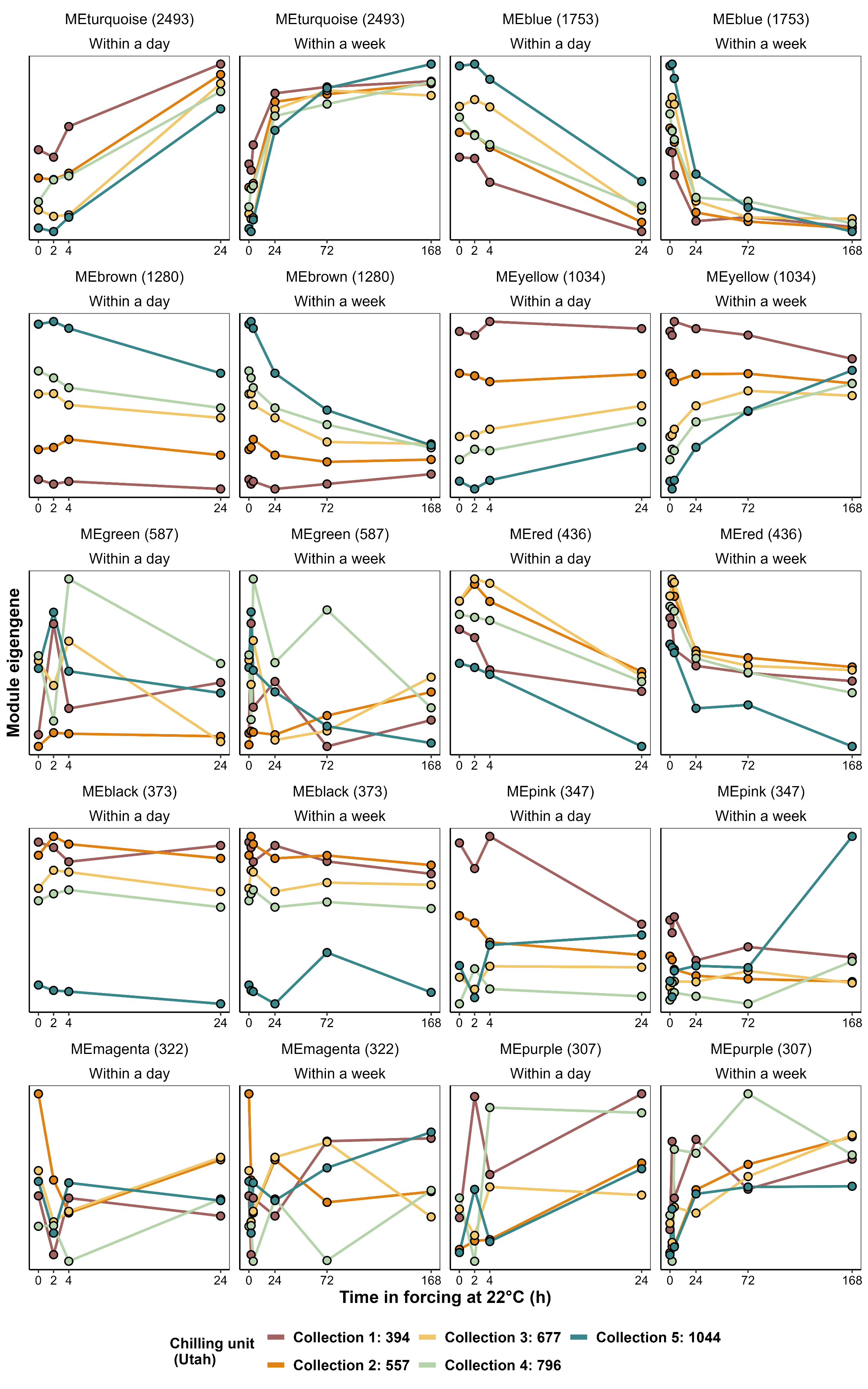

Figure S2. Expression pattern of the module eigengene (ME) of each gene co-expression module identified by weight gene co-expression network analysis (WGCNA)

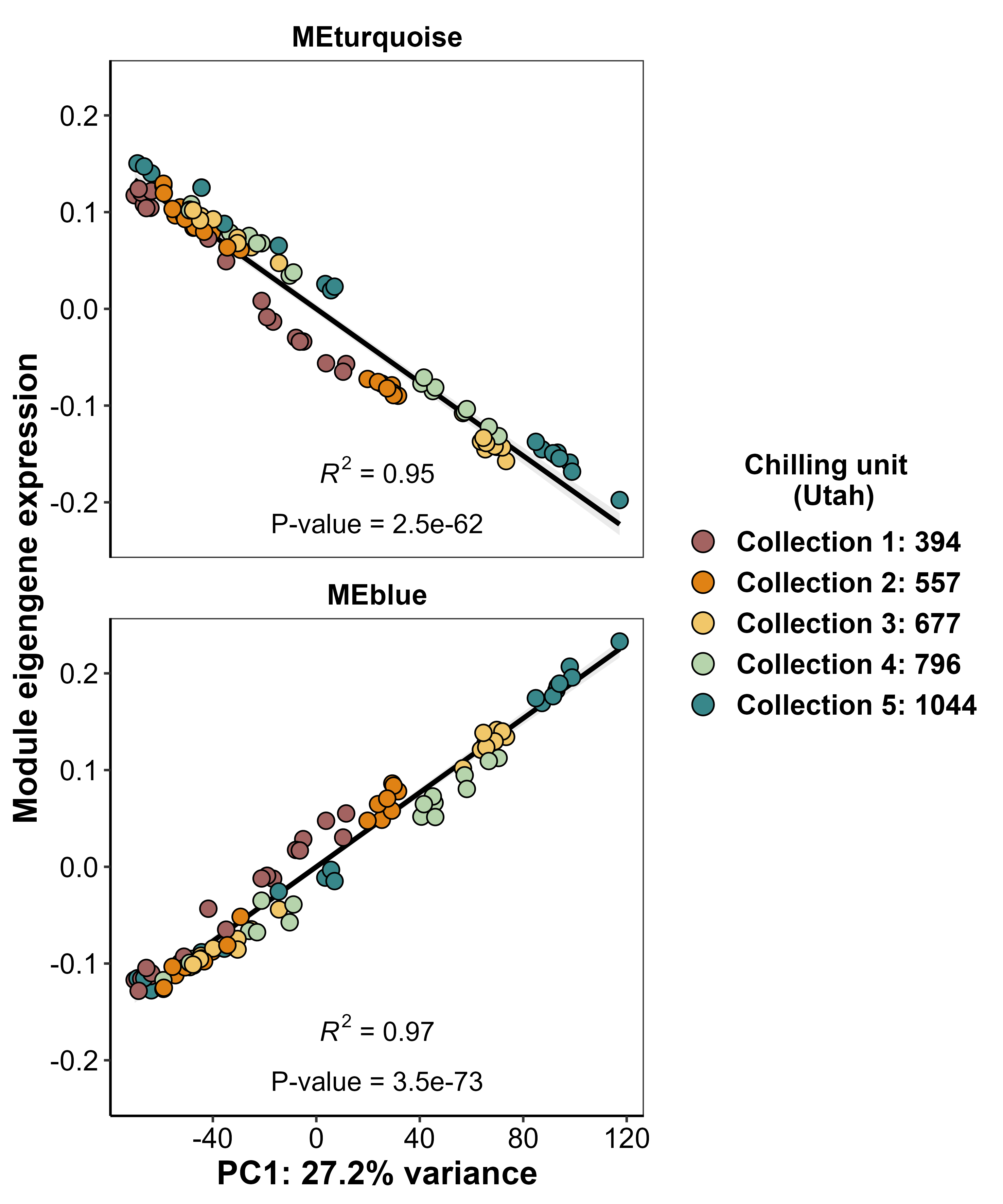

Figure S3. Correlation analysis of the MEs of gene co-expression modules ‘turquoise’ and ‘blue’ with PC1 from PCA.

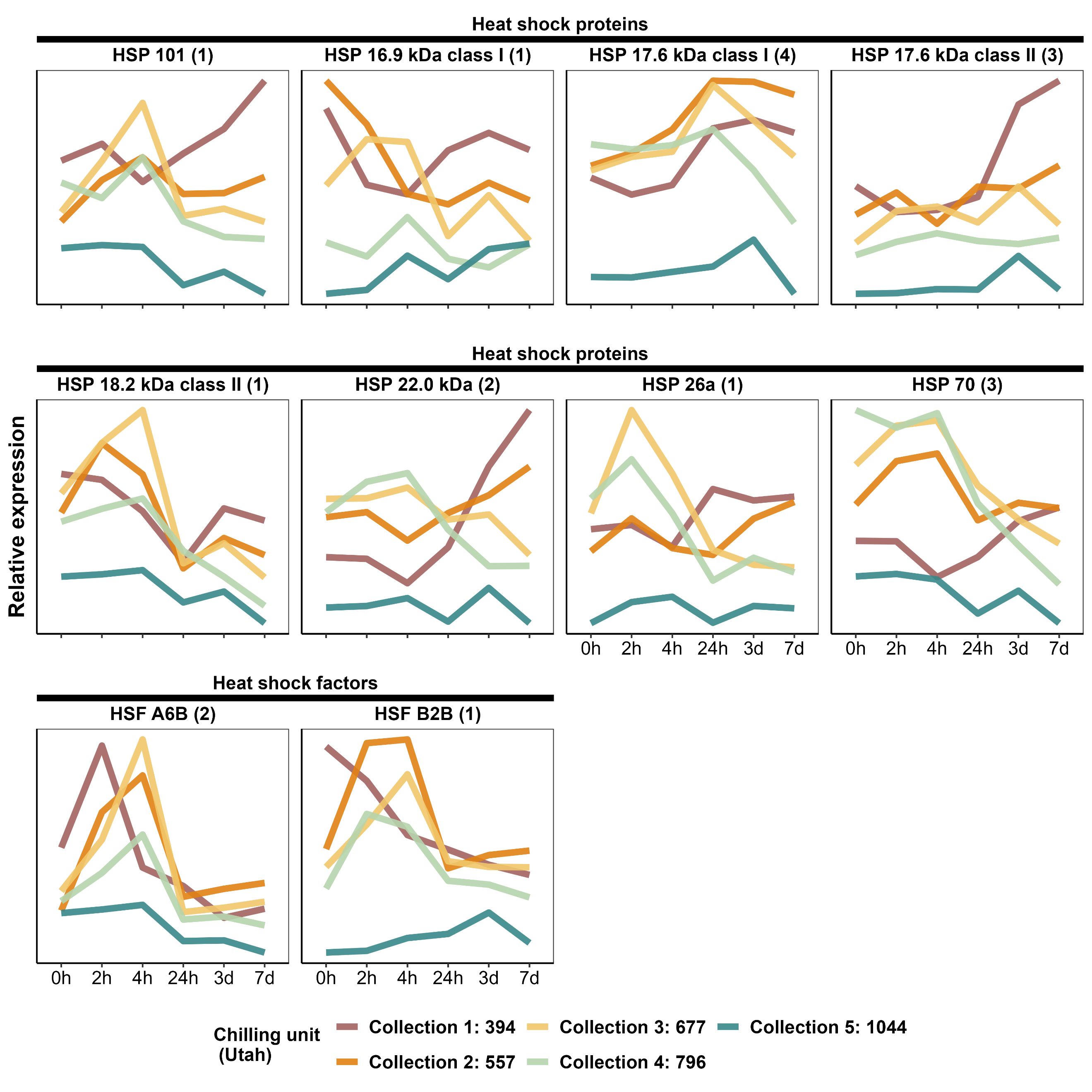

Figure S4. Expression of the heat shock protein (HSP) and heat shock transcription factor (HSF) genes significantly correlated with chilling. For clarity, expression values were summed for genes encoding the same protein. Numbers beside protein names indicate the number of gene copies included in each sum.

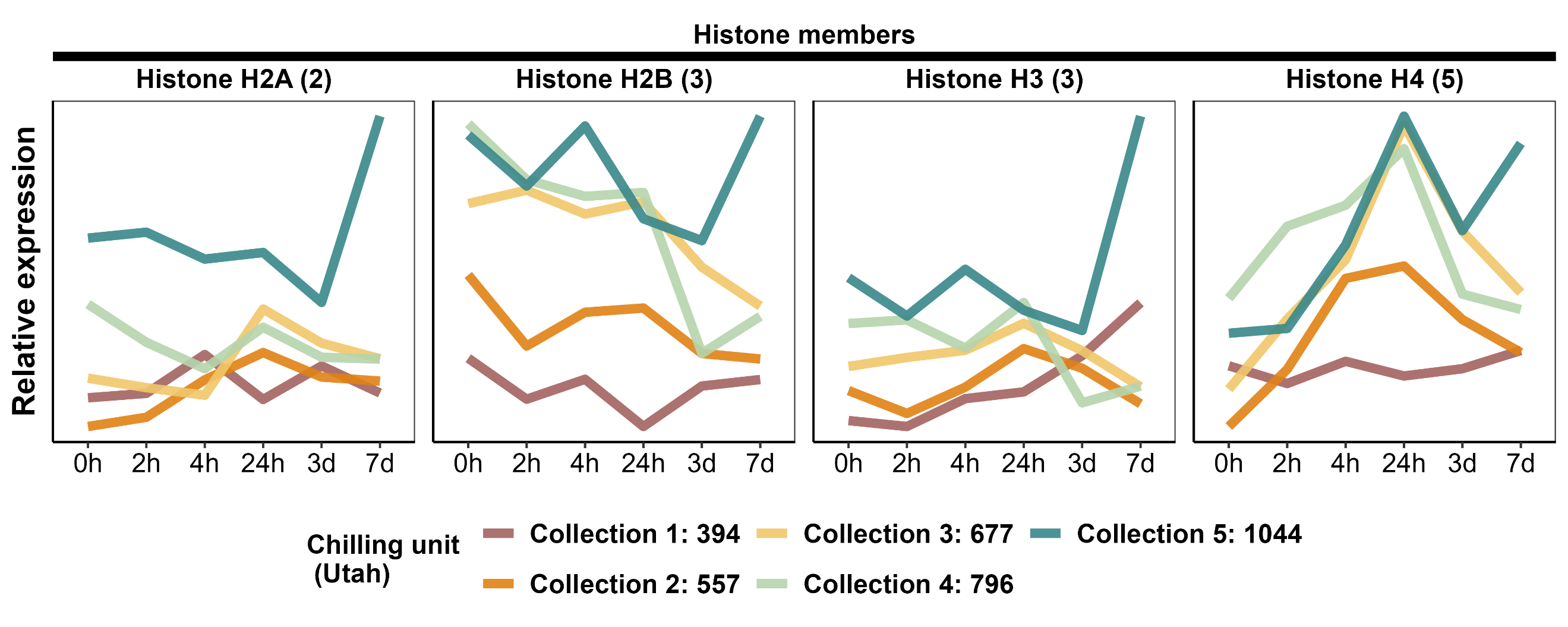

Figure S5. Expression of the histone octamer protein (H2A, H2B, H3 and H4) genes significantly correlated with chilling. For clarity, expression values were summed for genes encoding the same protein. Numbers beside protein names indicate the number of gene copies included in each sum.

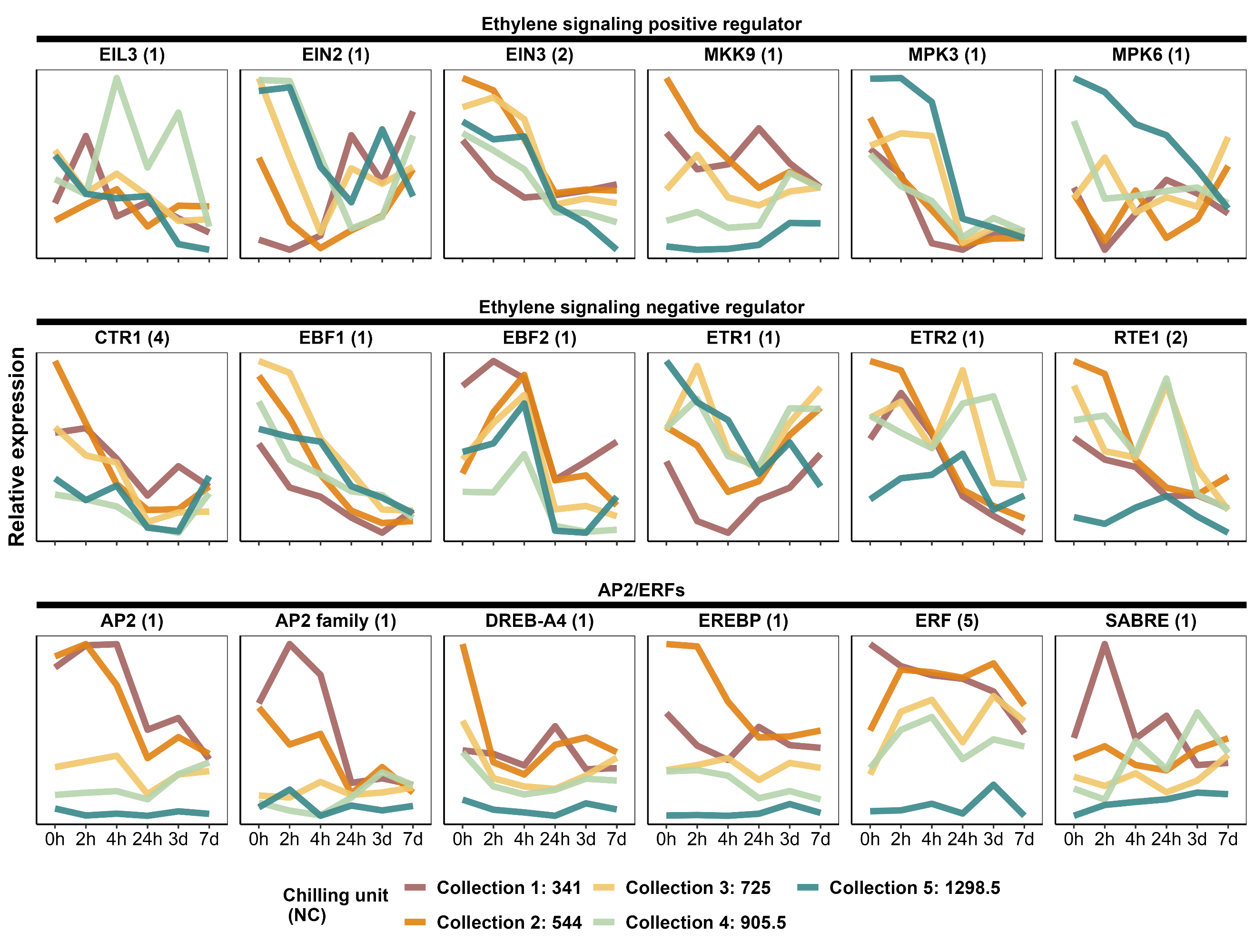

Figure S6. Expression of all the genes encoding ethylene signaling components that showed response to warm temperature exposure, chilling, or their interaction. For clarity, expression values were summed for genes encoding the same protein. Numbers beside protein names indicate the number of gene copies included in each sum.

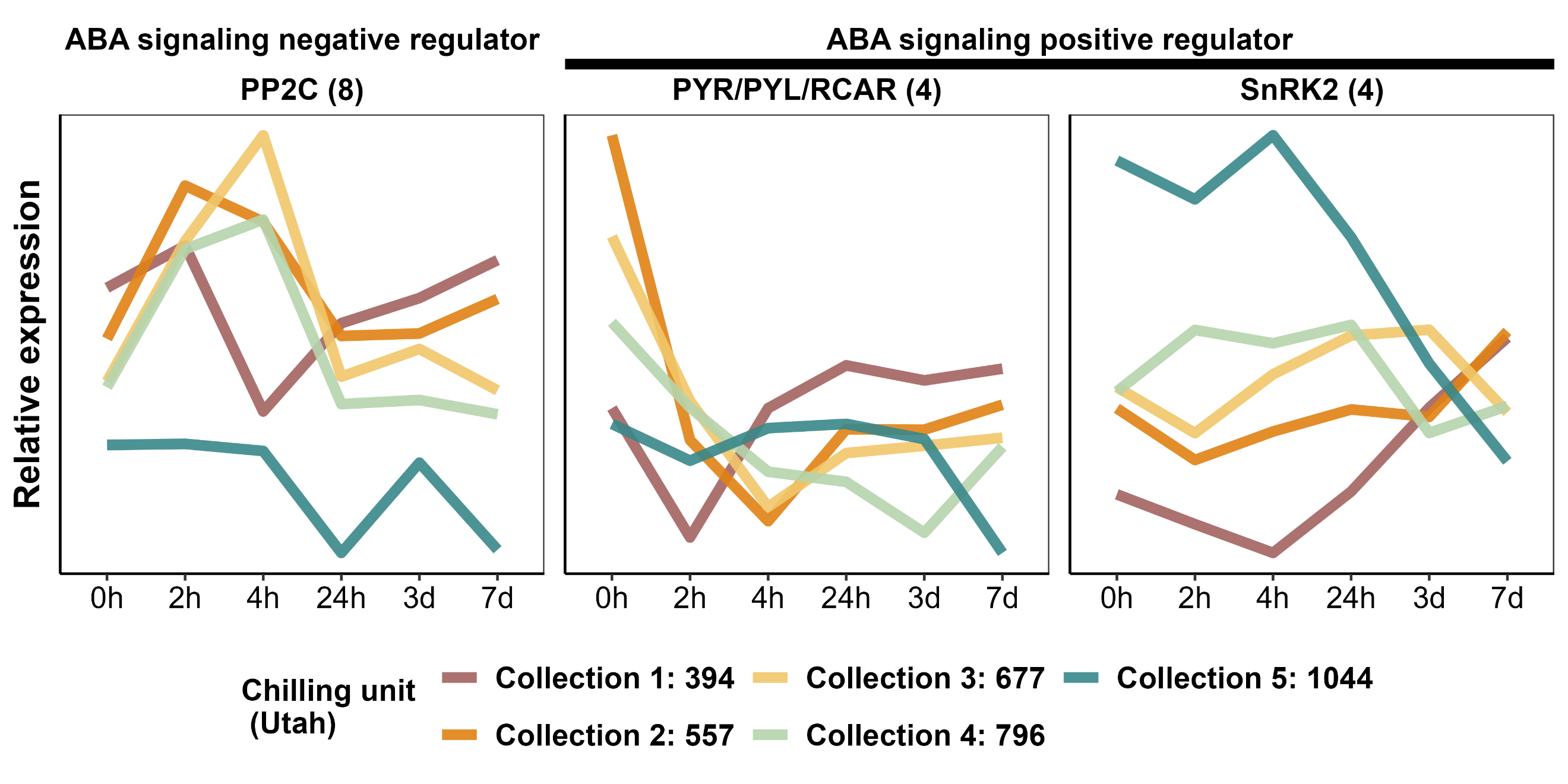

Figure S7. Expression of all the genes encoding three major ABA signaling components. For clarity, expression values were summed for genes encoding the same protein. Numbers beside protein names indicate the number of gene copies included in each sum.

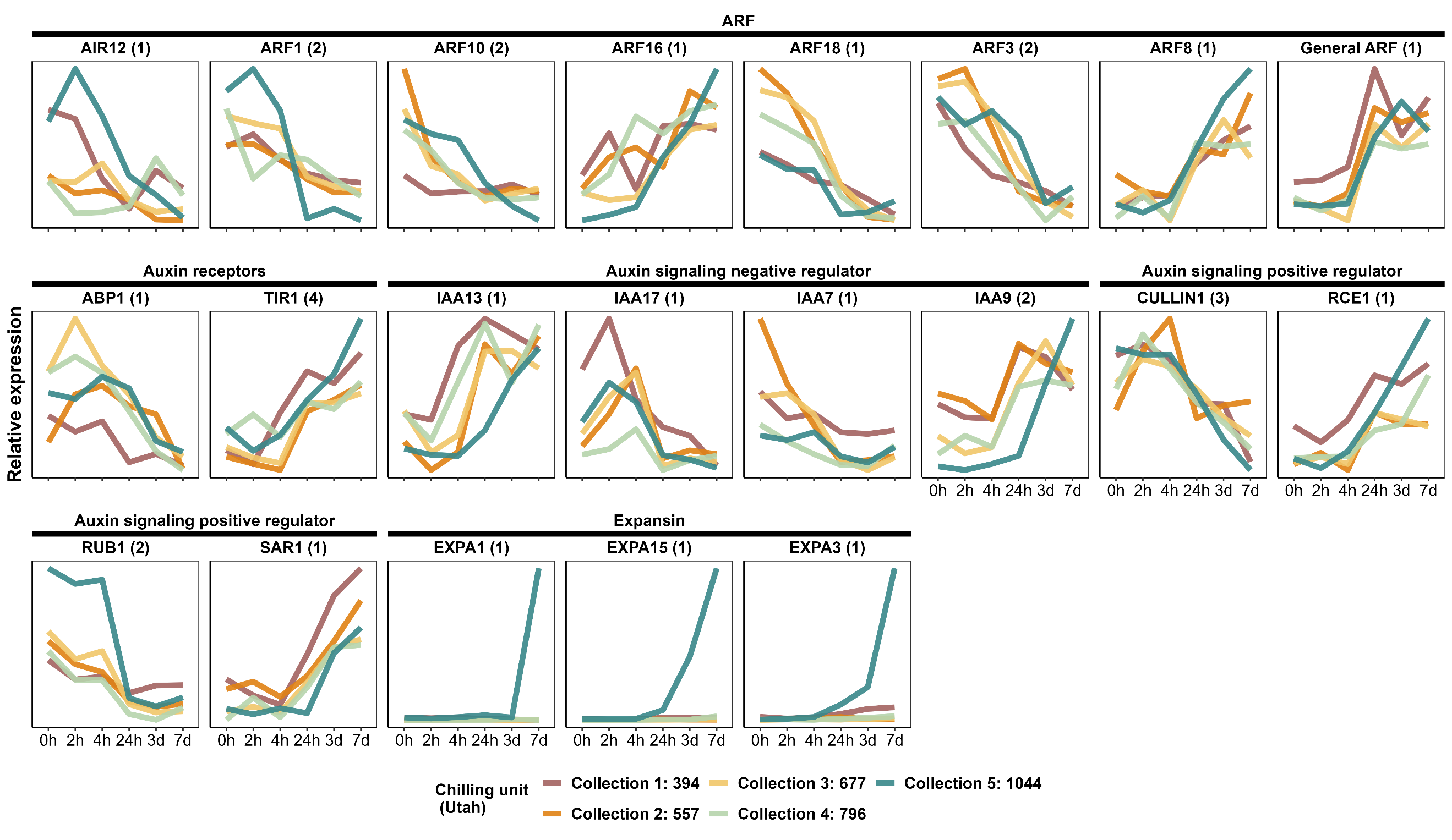

Figure S8. Expression of all the genes encoding auxin signaling components that showed response to warm temperature exposure, chilling, or their interaction. For clarity, expression values were summed for genes encoding the same protein. Numbers beside protein names indicate the number of gene copies included in each sum.

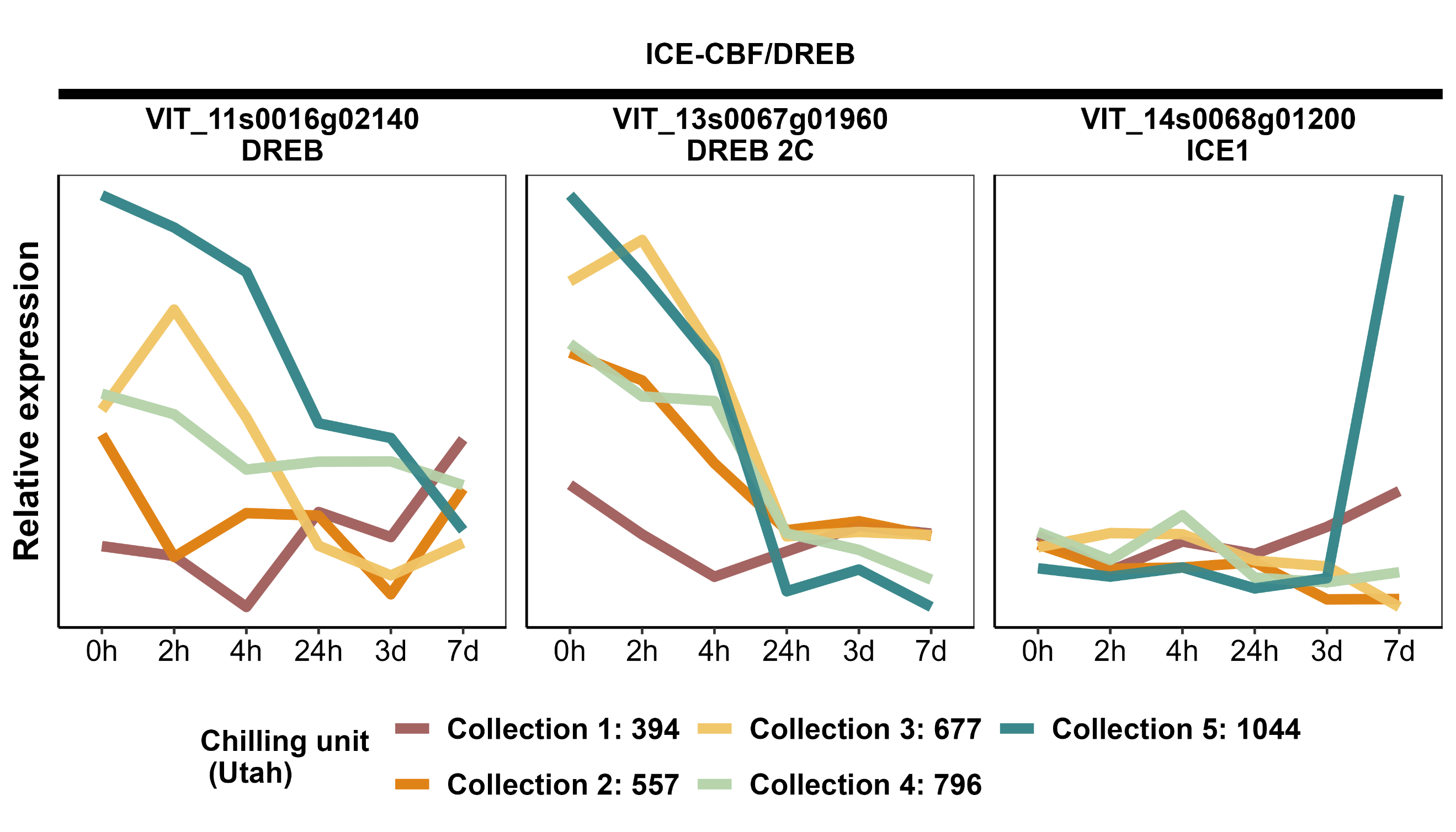

Figure S9. Expression of all genes with detectable expression encoding ICE or CBF/DREB.
